# Supply-driven evolution explains the locking-in of structures that open new evolutionary possibilities, such as higher hierarchies of organization

**DOI:** 10.1101/2022.07.18.500397

**Authors:** Julian Z. Xue, Leonid Chindelevich, Frédéric Guichard

**Author notes:** Email address:* (Julian Z. Xue, Leonid Chindelevich, Frédéric Guichard).

## Abstract

Many well-documented macro-evolutionary phenomena, such as increases in organization hierarchy, or sudden and episodic creation of new taxa, still challenge current evolutionary theories. Here we show a new mechanism that can explain them. We begin by showing how the order of mutations can alter evolutionary trajectories. We present a framework integrating both mutation and environmental biases and show that mutation bias can strongly outweigh any environmental bias, a regime we call “supply-driven” evolution. We then show how a common type of mutation bias, where detrimental mutations are more common than beneficial mutations, can drive the locking-in of structural hierarchies such as eukaryotism or multicellularity, independently from the environment. Finally, we generalize this result to show how any mutation (or sets of mutations) that creates the possibility of new phenotypes can persist for a very long period of time. That is, innovations that make possible a large range of new phenotypes can become frozen in time. By becoming frozen, these novel structures can no longer change, which means a range of phenotypes also become impossible. This opening and closing of phenotypic space is a new mechanism of macro-evolution.

## Introduction

One of the great and unresolved debates in evolutionary theory is the relationship between macro and micro-evolution. On the one hand, authors such as Dietrich (2009) provides an excellent overview for why macro-evolution should be an extension of micro-evolution, on the other hand, other authors such as Erwin (2000) are passionate and articulate about why macro-evolution is a separate process that cannot be distilled to micro-evolution.

Among the best arguments of the second school of thought are the many macroevolutionary phenomena that resist micro-evolutionary explanation. We might categorize them in four classes. The simplest class describe long term evolutionary trends (McShea, 1994) over macro-evolutionary time. For example, people (and scientists) have persistently, despite controversy in all its examples, thought about evolution as a long term progression in complexity and intelligence.

A better class of macro-evolutionary phenomena was described by Szathmary and Smith (2000); their examples of major transitions in evolution describe a steady increase in organizational hierarchy, such as from RNA molecules to prokaryotes, prokaryotes to eukaryotes, single-cell eukaryotes to multi-cellularity, and solitary individuals to eusociality.

The seeming irreversibility of major transitions in evolution relates this to a third class of phenomena, what has been described as “generative entrenchment” (Wimsatt, 1986; Schank and Wimsatt, 1986), that is, there are developmental events that lock-in in time. There are no members of the tetrapod class with more than four limbs, for example. Some developmental events in evolution simply matter more than other events, giving evolution the character of historical contingency (e.g. Powell (2009)) – a single such event in developmental pathways can alter evolutionary trajectory for all time for the lineage in which it occurs.

A fourth class of macro-evolutionary phenomena is proposed by the proponents of punctuated equilibrium. For example, Erwin (2000) says

Microevolution provides no satisfactory explanation for the extraordinary burst of novelty during the late Neoproterozic-Cambrian radiation (Valentine et al. 1999; Knoll and Carroll 1999), nor the rapid production of novel plant architectures associated with the origin of land plants during the Devonian (Kendrick and Crane 1997), followed by the origination of most major insect groups (Labandeira and Sepkoski 1993). Each burst was followed by relative quiescence, as the pace of morphological innovation fell.

These four classes of phenomena point clearly to the inadequacies of existing micro-evolutionary theory. However, those who critique micro-evolution have always struggled because modern macro-evolutionary theory does not offer better alternatives. Some of the macro-evolutionary mechanisms that have been proposed, such as species selection, cannot explain these phenomena. Others, such as the re-patterning of regulatory pathways (Erwin, 2000), are not (yet) rooted in mathematical theory. Without such theory, it is unclear how these explanations would exactly play out in the physical world, and how they would interplay with existing evolutionary theory.

We will here introduce a mechanism that can explain all four classes of macroevolutionary phenomena described above, and show that they may have a single root. We will show how this mechanism is entirely compatible with existing theories of micro-evolution, but is itself a firmly macro-evolutionary explanation, because it is a pattern that we can only see after many rounds of natural selection. Our work is rooted in ideas of mutation bias. We first claim that the order and frequency in which mutations occur can alter evolutionary trajectories. We then show how, in same cases, these biases can create long term trends in evolution that is not reversible by the environment. We next show how this implies that there is a special class of evolutionary innovations: those that create the possibility of new phenotypes. These evolutionary innovations can become “locked in” over evolutionary time. The arrival of these innovations, their locking-in, and the subsequent exploration of the new phenotype space they make possible create the phenomena of major evolutionary transitions, generative-entrenchment, and macro-evolutionary explosions.

### Mutation bias as an evolutionary driver in macro-evolution

For a long time, evolutionary biologists rejected the idea that mutation bias can be a significant force in evolution. This rejection is rooted in early models by Fisher that pitted mutation bias against natural selection. These models showed that when two alleles are in competition with each other, even a small amount of natural selection in favor of one allele easily overwhelms even large amounts of mutation bias in the opposite direction, especially when mutations are rare (Fisher, 1999).

The assumption Fisher made, however, was that both alleles were already available. However, under conditions where not all variations are immediately available, mutation bias is a fundamental driver of evolution. This is because selection can only work on that variation which is presented to it, meaning that whichever phenotype arrives first gets first chance at success (Fontana and Buss, 1994; Stoltzfus, 2006b; Stoltzfus and Yampolsky, 2009; Stoltzfus, 1999; Yampolsky and Stoltzfus, 2001; Schaper and Louis, 2014)

A similar framework comes out of quantitative genetics (Fig 1). Depending on the *G* matrix, which determines frequency of different mutations (in this case, an elliptical or circular mutation space), the population will discover a different fitness hill. Mutation bias can thus become a powerful force that dictates the direction of the evolution of a trait (Stoltzfus, 2006a).

**Figure 1:**
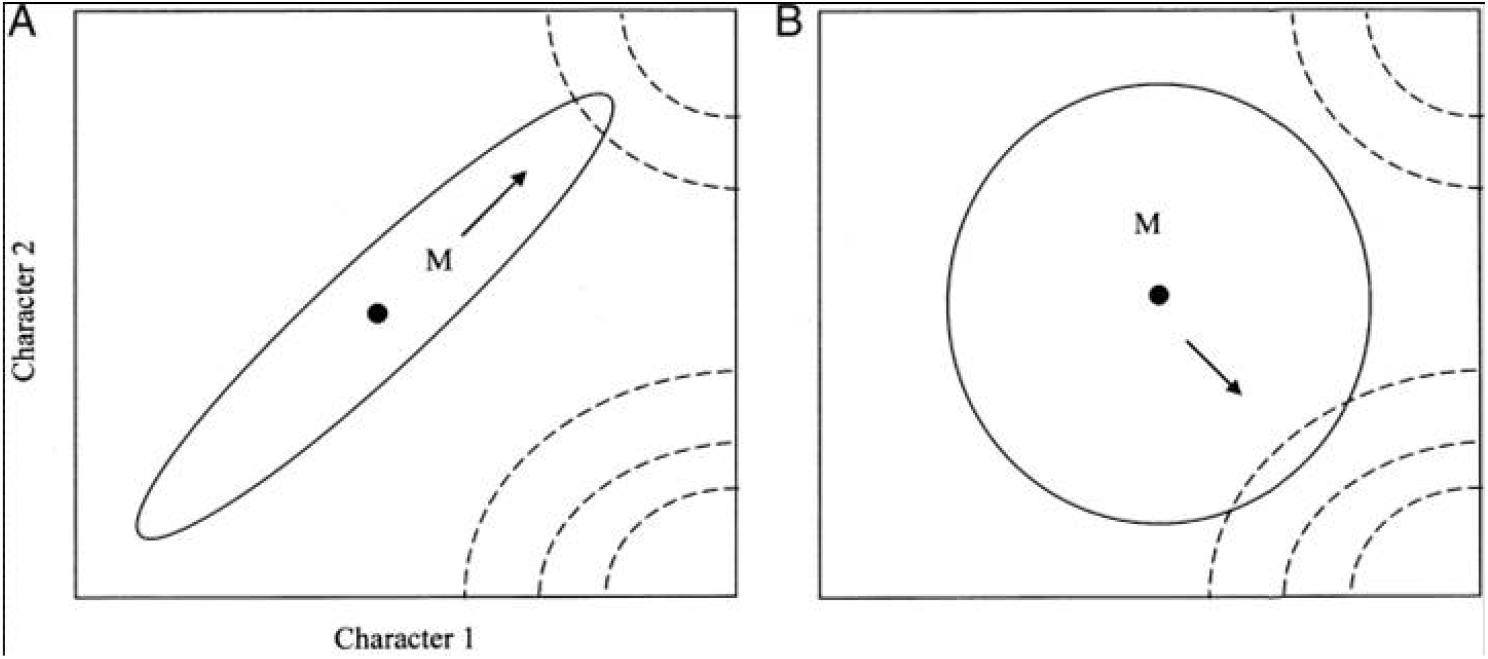
Mutation bias in quantitative genetics with multiple traits: there are two fitness hills near the population. Depending on how it mutates (ellipse vs. circle), it will choose one over another. Adapted from Arthur (2004)

Constraints on the availability of mutations have roots in developmental biology with well-documented constraints on the development of organisms imposing limits on - and thus biasing - genetic variation (Alberch and Gale, 1985; Klingenberg, 2005; Arthur, 2002; Müller and Newman, 2003; Hall et al., 2004). Theoretical frameworks inspired by quantitative genetics have integrated verbal arguments about mutation bias and experimental observations from developmental biology (Rice, 1990, 1998, 2012; Klingenberg, 2010). These approaches require measuring and making assumptions about the shape of multivariate landscapes, such as the trait variance-covariance (*G*) matrix, as well as the fitness landscape over many traits – never an easy problem. The complexity of these models also make intuitive insight difficult.

Alternative approaches include a simpler but crude model for thinking about mutation bias (Xue et al., 2015) based on the concept whereby two organisms with same trait value do not necessarily have the same fitness. As observed by Arthur (2001), changes in a trait value such as body length can be achieved via different developmental mechanisms such as increased cell size or cell proliferation. As another example, there are many ways for different organisms to have the same complexity level, no matter how complexity is measured. In this context, the relationship between trait value and fitness becomes probabilistic.

We consider mutation bias to be the phenomenon that different trait values can be introduced by mutation with unequal probabilities. Combining this with the idea that a given trait value can be produce by different mutations is the pivotal insight for our study.

#### Mathematical framework

Let us consider a homogenous haploid population with fitness *ω*\* and a trait with value *ϕ*\*. Let a mutant *m* be defined by *s*_*m*_, *u*_*m*_ where *s*_*m*_ = *ω*_*m*_ − *ω*\*, the difference between the mutant and resident fitnesses, and *u*_*m*_ = *ϕ*_*m*_ − *ϕ*\*, the difference between their trait values. A mutant-fitness distribution, ζ(*s, u*), can now be defined over all mutants of the population, which is a bivariate probability distribution over *s* and *u*. When a mutant enters the population, its fitness and trait value are determined by choosing from ζ. Ordinary evolution then takes place to determine the success of this mutant. Note how, in this framework, mutants with the same trait value *ϕ*_*m*_ can have very different fitnesses, drawn from the distribution ζ(*s, u* = *u*_*m*_) (see Figure 2).

**Figure 2.**
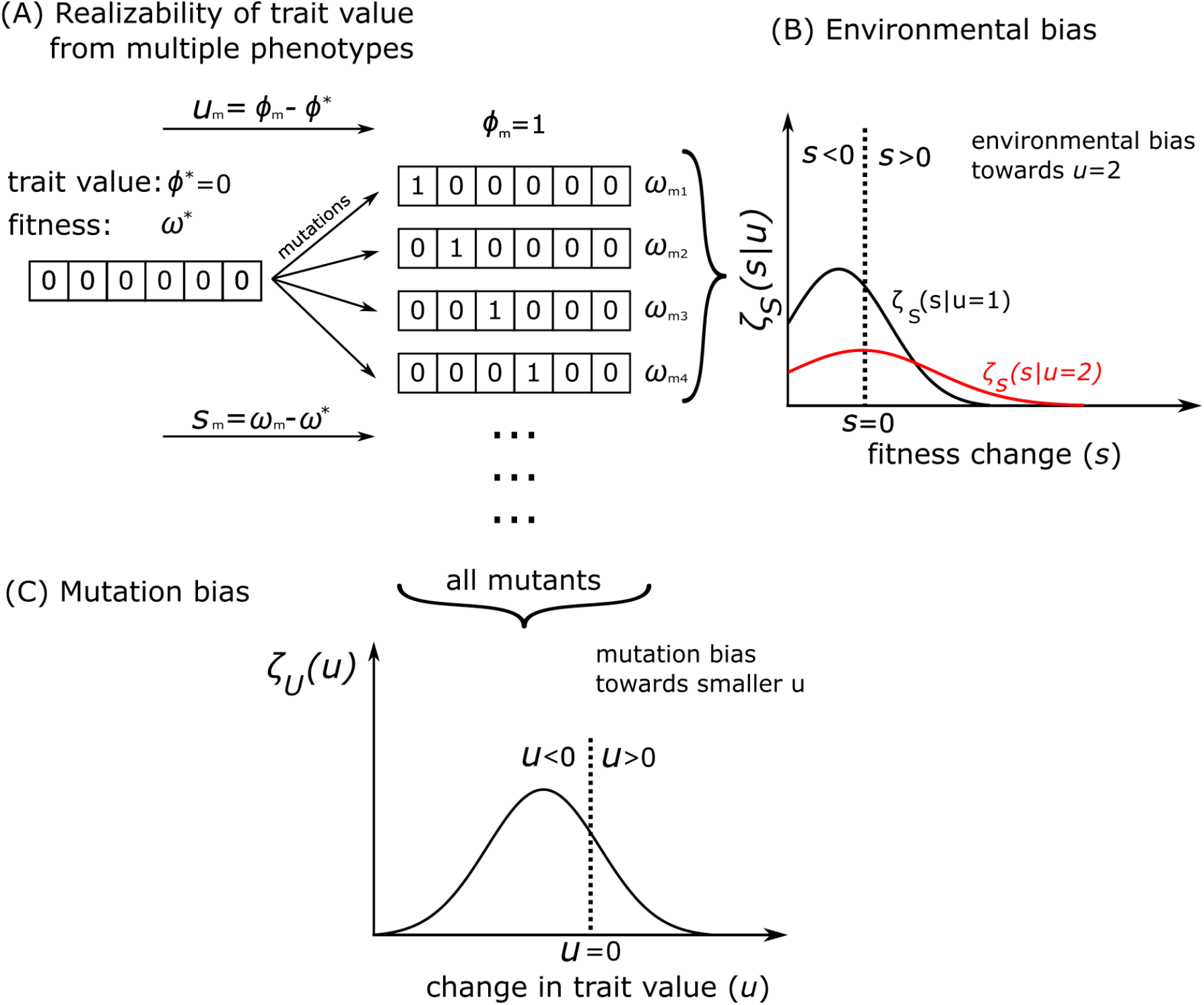
This figure shows how the NK model we use for our agent-based model implements environmental and mutation bias. We only show the values of each component. The fitness table and the links between components are not shown for clarity. On the top left, we have the wildtype, whose components are all 0’s. It has fitness *ω*\* and trait value *ϕ*\* = 0. To its right are some of its possible mutants. These mutants all have the trait value *ϕ*_*m*_ = 1 (so *u* = 1 also), but they are different phenotypes and have different fitnesses, which form a distribution over *s* (the difference between mutant and wildtype fitnesses), shown on the right: ζ_*S*_(*s*|*u* = 1). If we do not take the set of all mutants *ϕ*_*m*_ = 1, and instead took the set of all mutants *ϕ*_*m*_ = 2 instead, we would have a different fitness distribution, ζ_*S*_(*s*|*u* = 2) (illustrated in red). The fact that this distribution is right-skewed compared to ζ_*S*_(*s*|*u* = 1) means that the environment favors *u* = 2. If we took the set of all mutants and simply plotted them all (regardless of their fitness) according to their *u* value, we would have the mutation bias distribution, ζ_*U*_ (*u*) (the bottom plot of this figure). Here, we see there is a mutation bias towards smaller *u*.

We will work within a classic adaptationist scenario, where there is strong selection and weak mutation (for a categorization of the different regimes of evolution, see Sniegowski and Gerrish, 2010). In this case, mutations are rare. When a beneficial one comes along, it can sweep through the population if it escapes drift. In the ideal case, the only driver of evolution in this scenario are selective sweeps, which happen on a timescale that is short compared with the timescale of the generation of beneficial mutants. To simplify our analytical model, we will assume that detrimental mutations are evolutionarily inconsequential, that there is no clonal interference (that is, there is only one adaptive mutation at a time – see Kim and Orr (2005)), and that improvements in fitness are small and incremental. We will see with agent-based models that relaxing these assumptions does not change the final results.

Since improvements in fitness are small, the probability of a mutant with fitness advantage *s* sweeping the population is *αs*, where *α* is a constant that depends on the details of evolution (*α* = 2 for Poisson distributed offspring (Haldane, 1927), *α* = 1 for a Moran process (Nowak, 2006), *α* ≈ 2.8 for binary fission – see the discussion in Johnson and Gerrish (2002)). Thus, the probability distribution that a mutant will sweep the population and change the trait value by *u* is a function of *u*:

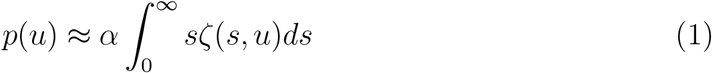

We integrate over *s* from 0 to ∞ because we only care about the higher fitness mutants. The probability *p*(*u*) does not normalize to 1 over all *u*, the probability that there is no selective sweep makes up the rest of the weight.

Of course, ζ(*s, u*) may change over time, but for now we will consider a single selective sweep before taking a long term view.

To simplify this equation and to understand it better, we first can separate out the contribution of mutation bias to evolution, We can split ζ up according to its conditional probability distribution, using the standard notation where *U* and *S* are random variables for *u* and *s*:

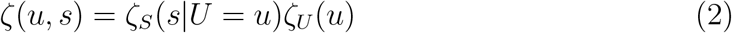

Here, ζ_*U*_ (*u*) is the marginal density of mutants that change trait value by *u*, while ζ_*S*_(*s*|*U* = *u*) is the fitness distribution of all mutants with trait value difference *u*. In essence, ζ_*U*_ (*u*) is the mutation distribution: it tells us how likely a mutant with particular *u* is likely to arrive. On the other hand, the environment is captured by ζ_*S*_(*s*|*U* = *u*): for a given trait value, the environment determines the fitness distribution of all mutants with that trait value. For example, if the environment becomes more favorable for a trait value *u*, the mean of the distribution ζ_*S*_(*s*|*U* = *u*) may increase. Of course, the environment can change ζ_*S*_(*s*|*U* = *u*) in many other ways, such as increase its variance or skew, which can also qualitatively alter evolution.

Next, we split ζ_*S*_(*s*|*U* = *u*) into two parts, the detrimental and the beneficial parts

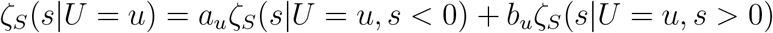

where

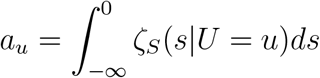

and

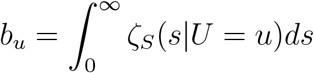

That is, *a*_*u*_ is the probability that a mutation that changed the trait value by *u* is detrimental, and *b*_*u*_ is the probability that a mutation that changed the trait value by *u* is beneficial. In this case, equation (1) becomes

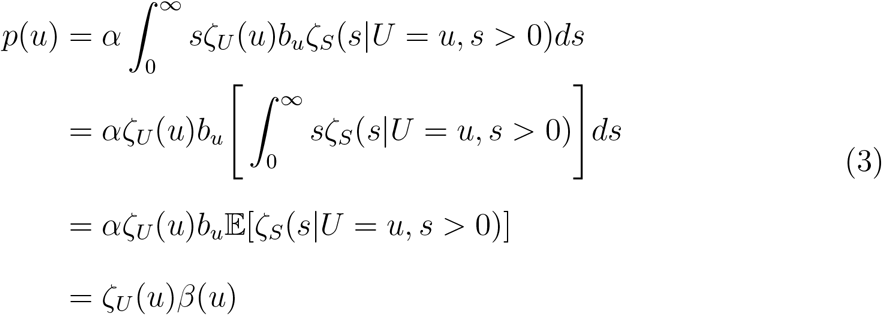

Here, *β*(*u*) = *αb*_*u*_𝔼[ζ_*S*_(*s*|*U* = *u, s >* 0)] can be called the “degree of benefit” of *u*, which is the probability that a mutation that changed the trait value by *u* is beneficial, *b*_*u*_, multiplied by the size of the expected fitness advantage, 𝔼[ζ_*S*_(*s*|*U* = *u, s >* 0)], multiplied by *α*. What *β* measures is the probability that a mutation with trait value *u* will selectively sweep – that is, the mutation will both be beneficial as well as escape drift. Detrimental mutations drops out of the equation because the integration on *s* is only from 0 to ∞, that is, in this framework only beneficial mutations will alter the evolutionary trajectory.

Equation (3) defines the probability that the population trait value will change by *u* through the introduction of the next mutant as the likelihood of this mutant appearing, ζ_*U*_, multiplied by the degree of benefit of this change, and a factor *α* that depends on the exact nature of the evolutionary process as described earlier. In this equation, the mutation distribution plays an equal and symmetric role to environmental selection. A trait value with half the benefit of another trait value can make up for it with double the mutation bias.

We can integrate through all trait values, *u*, to arrive at the expected change in trait value through the introduction of one mutant:

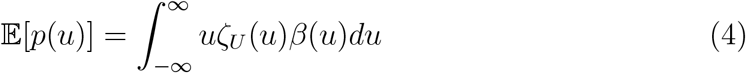

To recap, the above equation holds in regimes where the only evolutionary events are selective sweeps by beneficial mutations, where beneficial mutations are rare and the fitness effect of beneficial mutations are small. In this formulation, selection affects evolution by changing *β*(*u*), while mutation bias is captured by ζ_*U*_ (*u*). The last equation holds for the introduction of a single mutant. As time goes on, both ζ_*U*_ and *β* may change: the former as the structure of the organism changes, thus changing the sort of mutants it produces, and the latter will change as the environment changes.

#### Prediction of trait change in an agent-based model

We can first apply equation (1) to predict how a trait evolves in agent-based simulations where ζ_*U*_ (*u*) and *β*(*u*) are known or can be measured. To demonstrate this, we adopt an agent-based model that was previously used to show that mutation bias can direct the evolution of a trait (Stoltzfus, 2006a), even against environment selection (Xue et al., 2015).

The model is an agent-based NK-model (Kauffman and Levin, 1987) where agents are strings made from elements of 0’s and 1’s. There are interactions among these elements that create a rugged fitness landscape. Mutations flip 0’s to 1’s and viceversa. The trait we measure is the sum of all the elements. The environmental bias is defined in such a way that the larger this sum, the more likely that the agent is fit. There is a mutation bias in the opposite direction, however, in the sense that mutations from 1 to 0 is more common than mutations from 0 to 1 (Fig 2, see Appendix A for details).

In this model, ζ_*U*_ is known because it corresponds to how often mutations change 0 to 1 or vice versa. *β*(*u*) measurable because it is a dynamic quality determined by how close the population is to a fitness local optimum. *β*(*u*) is affected by many factors: the height of the fitness optimum, the rate of change in the environment, and by the mutational load, among other things.

*β*(*u*) changes over time. Since *β*(*u*) measures the degree of benefit provided by a trait value, it will change with the environment – as a population gets close to the optimum of an environment, *β*(*u*) ≈ 0 for all *u*, since there are much fewer mutants with improved fitness. If the environment changes constantly at a regular pace, however, we can predict *β*(*u*) to remain relatively constant over time. This is what happens in this model: with a single measurement of *β*(*u*), we can make good predictions for the entire time period (Figure 3). In these cases, then, the actual long term rates and directions of a trait’s evolution can be known if we can obtain a measurements for *β*(*u*).

**Figure 3:**
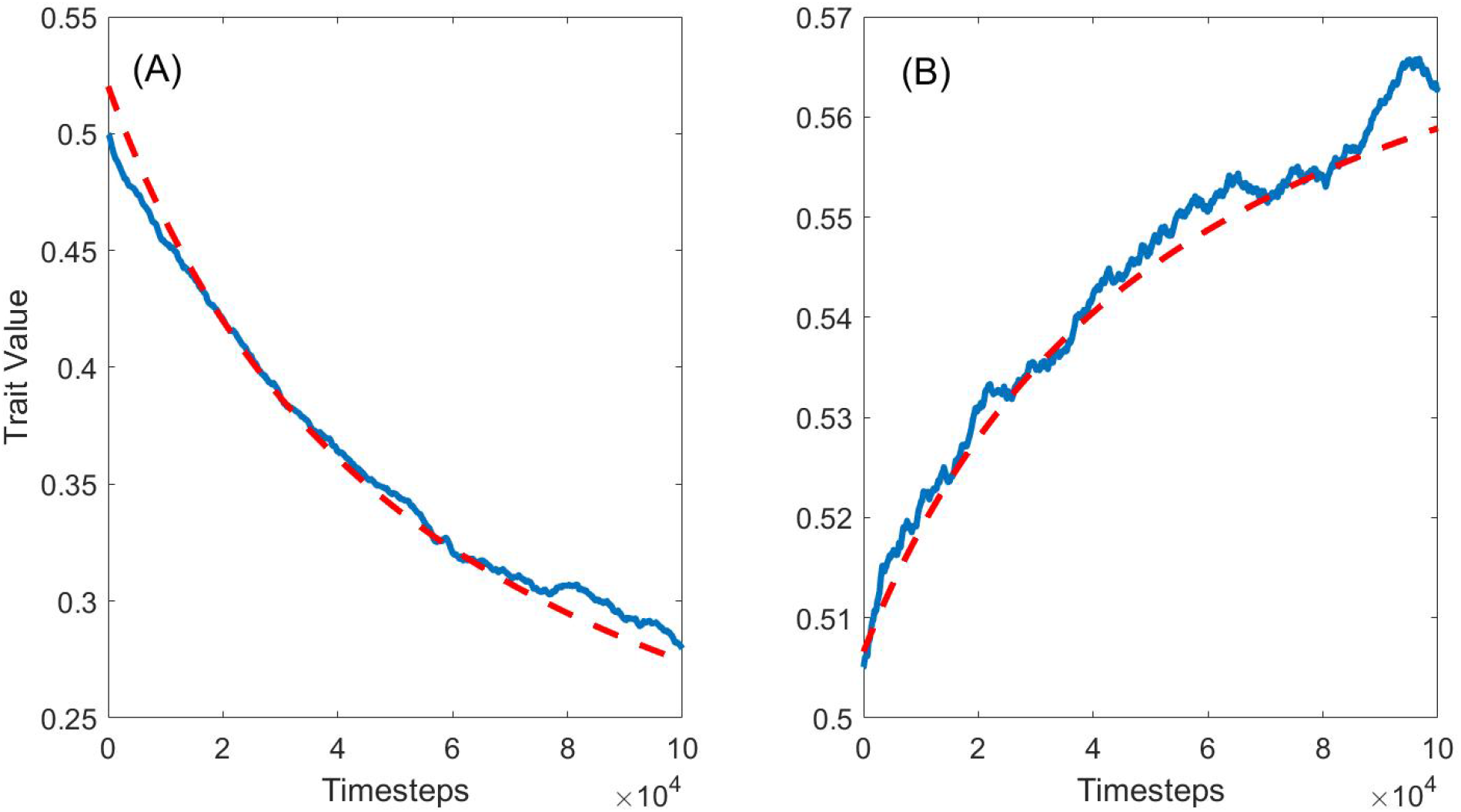
The blue line is the average trait evolution for 200 model runs, the red dashed line is the predicted evolution by equation (4). For the set of runs in panel A, mutation bias pressures the trait value down while the environment is biased to larger trait values. The environment changes over time. Panel B is identical, but without mutation bias, allowing environmental bias to push the trait value upwards. In both plots, *β*(*u*) was measured at *t* = 2*e*5, giving the model time to find an equilibrium between environmental change and mutation-selection. After that, *β*(*u*) stayed relatively stable over time, so a single measurement allowed us to make good (although imperfect) predictions over the entire simulation both backward and forward through time. Measuring *β*(*u*) at *t* = 0 was not as helpful, since the population has not settled in an equilibrium with the environment.

### The supply-driven evolution of structural hierarchies

We will now study scenarios where the shape of ζ_*U*_ (*u*) introduces a mutation bias that is unlikely to be reversed by selection from the environment. The evolutionary directions of associated traits are determined by the supply of mutation and can be called “supply-driven evolution”. While the speed of evolution can be affected by the environment, its direction will not. We will now study how such a scenario can explain the evolution of structural hierarchies.

The evolution of structural hierarchies capture major transitions in evolution (Szathmary and Smith, 2000) including transition from the RNA world (Orgel, 2003) to the first cells, from prokaroytes to eukaryotes (Lang et al., 1999), from unicellular to multicellular eukaryotes (Bonner, 1998), and from multicellular organisms to eusociality, such as ants or naked mole-rats, and perhaps humans (Wilson and Hölldobler, 2005). What is most interesting about these major transitions is their irreversibility. As far as we know, there is no free-living unicellular organism that had an animal, plant, or fungi as its ancestor. Nearly all animals get cancer, but no cancer cell is known to survive independently of the host other than in highly controlled lab conditions. At the very limit, there are cancers in dogs, clams, and Tasmanian Devils that can infectiously spread, but even they are not known to live autonomously (Metzger et al., 2015; McCallum, 2008). There are a few exception to this irreversibility, since there are solitary organisms that had eusocial ancestors (Wcislo and Danforth, 1997) and there are eukaryotes that have lost their mitochondria (Clark and Roger, 1995; Horner and Embley, 2001), although they usually retain a remnant (Tovar et al., 1999, 2003; Tachezy et al., 2001; Hampl and Simpson, 2007).

Although there have been explanations for the origin of particular increases in hierarchy, we note that there is no consensus on a fitness based theory for why the increase in hierarchy should persist. Prokaryotes are, by almost any measure, the most successful kingdom on the planet. The main other type of explanation is drift to fixation of deleterious allele in each of the components organisms (e.g. Moran (2003)), rendering the hierarchy obligatory. In the next section, we show how this will occur with high probability even when there is no neutral or deleterious drift to fixation and only selection takes place.

#### The mutation distribution drives the locking-in of hierarchies

When two component organisms live in such close synchrony that they form one unit of selection – that is, they share a measure of fitness, and mutations in each affect this joint fitness, then mutations on either component have two effects: one on the fitness of the joint ensemble, and another on the fitness of individual components, should they separate from the whole. To see how locking-in first use of ‘locking-in’, define happens, we simply have to show a scenario of how the fitness of each independent component would steadily degrade over time. Components may begin as autonomous organisms, but become fully dependent on the whole over time and unlikely to have a viable fitness when severed from the whole.

We first define the fitness of the component organism as a trait with value *ϕ*. We assume mutation bias leads to more frequent detrimental mutations compared to beneficial mutations (Eyre-Walker et al., 2006; Eyre-Walker and Keightley, 2007; Monroe et al., 2022). We then define *ω* as the fitness of the joint ensemble. We now have *u* = Δ*ϕ* being the change in fitness of the components, should they become separated, while *s* = Δ*ω* is the change in fitness of the joint ensemble, which is what selection is immediately acting on.

Since the trait variation *u* is also a change in the fitness of individual components, we can use Fisher’s framework (see, for example, Orr (2006)) considering organisms to be points in a high dimensional space, where each dimension represents a trait. Mutations occur within a mutational volume (isn’t it strange to talk about a sphere in a space with an arbitrary number of dimensions?) around the organism, and the environmental optimum is another point in this space. Using this framework, theory and experiments show that detrimental mutation always outnumber beneficial ones, this relation becoming stronger as the organism approaches the environment optimum (Eyre-Walker et al., 2006). Assuming this inequality between detrimental and beneficial mutations and its relationship with the environmental optimum, we have:

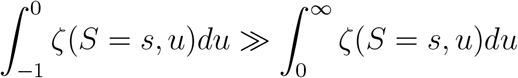

as we approach the environmental optimum. Since *u* is a change in fitness, its range is [−1, ∞].

However, this implies that

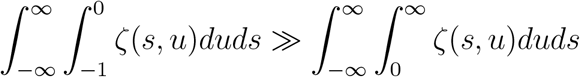

Switching the order of integration,

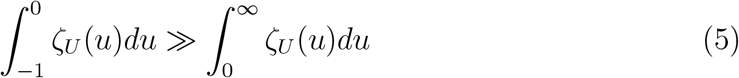

From this maybe be more specific about what eq. 5 means and why it can lead to the next step/prediction, our aim to show that 𝔼[*p*(*u*)] < 0, that is, we expect the fitness of the component organisms to degrade at each selective sweep, such that over time they can no longer live autonomously.

In order to achieve this, we must define the relationship between trait and fitness variations. It has first been shown that the magnitude of detrimental mutations among the component organisms are, on average, larger than the magnitude of beneficial mutations (Eyre-Walker and Keightley, 2007). That is, the average detrimental mutation is more detrimental than the average beneficial mutation is beneficial, even if detrimental mutations have a lower bound of −1 while beneficial mutations have no upper bound. This relationship means that:

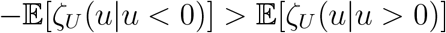

Since

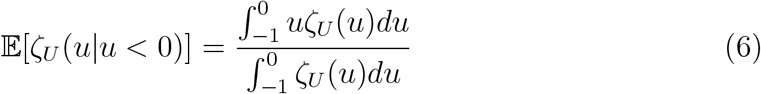

And similarly for 𝔼[ζ_*U*_ (*u*|*u >* 0)], multiplying the equations (5) and (6) gives us

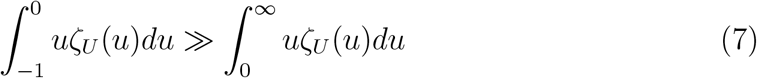

We also need to bound values of *β*(*u*). Given the fact that *β*(*u*) measures the degree of benefit (the probability that trait value *u* would be beneficial, multiplied by the expected degree of fitness increase if it were beneficial), *β*(*u*) must be bounded from above: that is, there are some trait value *u* for which *β*(*u*) is maximized. Of course, *β*(*u*) is also bounded below by 0. Thus, *β*(*u*) is also bounded above and below for any range of *u. β*(*u*) therefore has a minimum 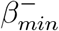 for −1 *< u <* 0. Similarly, *β*(*u*) has a maximum 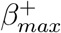 for *u >* 0. From these constraints we make the assumption that and 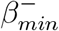 are such that:

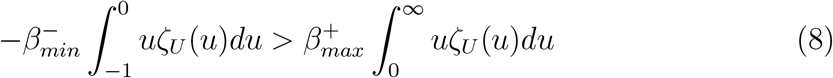

This is an important assumption. It means that 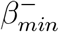 is not much smaller than 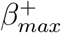. This has a neat biological interpretation: mutations that are detrimental to the fitness of the component organism, that is, decreases *u*, must still have a significant chance to increase the fitness of the whole organism, *s*. Equation (8) is a sufficient rather than necessary condition, and it is actually quite stringent, since it implies 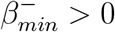, that is, mutation of any detrimental effect to the joint organism don’t you mean ‘to the component organisms?’ has a chance to improve the joint organism.

Once we have made these assumptions, equation (8) implies:

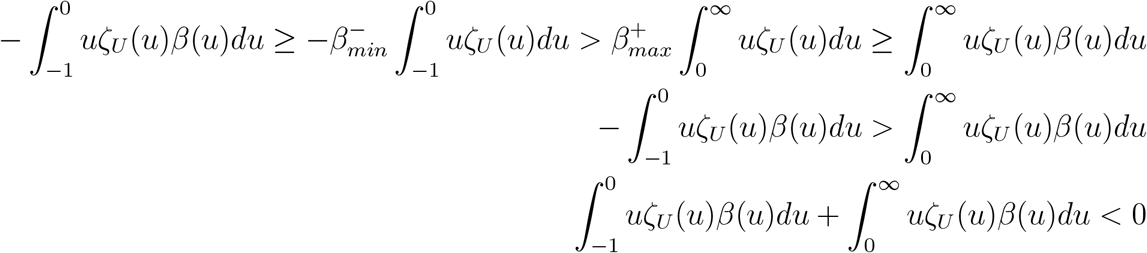

The last expression implies

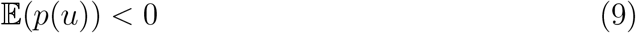

This means that, under these assumptions, we can expect the fitness of the component organisms to degrade at each selective sweep until there are more similar levels of detrimental mutations to beneficial mutations I don’t understand the previous statement: ‘more similar levels’?. At this point, the fitness of the component organisms is likely so low that that the joint organism is very unlikely to be separated into its component organisms, thus locking-in the new hierarchy of organization.

#### Evolutionary lock-in: an agent-based model

We apply the above theory to a NK model, so agents are strings made from elements of 1’s and 0’s that interact with each other. However, mutations does not change the elements; instead, they change interactions. Each mutation causes a number of elements to interact with different elements than before. We also have a rarer class of mutations: occasionally an agent will be joined with another agent, to form a much longer string. We call this a joining mutation. Initially, the elements of the two agents have only a few links with each other, but with mutations they eventually can be more heavily linked to each other. Lastly, long agents can also be cleaved in half to form two individual agents. The interactions that were present between the elements of the two agents when they were a single organism will then be lost.

The environment is constructed such that the longer the agent, the less likely it is to be fit. This is a conservative assumption, to show that joining mutations can be locked-in even when the environment is biased against it. There is a 2% penalty to fitness each time the length of the organism doubles. Increasing this penalty increases the amount of time before evolutionary lock-in holds; the system simply has to wait for a joining mutation that increases the fitness for the resulting joined organism, despite its increased length. This is possible because there are different ways for agents to join together, some of which cause an increase in fitness that can overcome the 2% penalty. For details on the model, see Appendix B.

Time series of organism length over multiple simulations (Figure 4a) illustrate how many joining mutations almost immediately reverse themselves, but some persist. This persistence means that the number of hierarchies rise over time, despite an explicit fitness penalty for doing so. Focusing on a single simulation where locking-in is observed (Figure 4b), we can show that the hierarchy rises when two components become increasingly enmeshed with each other as a larger unit as measured by the number of links between the first and second halves of the joint organism (Figure 4c). This is a measure of how much effect a cleaving mutation will have on the organism. This steadily increases with time, which means that a cleaving mutation will have increasing effect on phenotype over time, reducing its chances of being beneficial. On the other hand, *u*, the fitness of the component organisms, steadily declines over time (Figure 4d-f). We can see how the component organisms are becoming less and less fit and how much more frequent detrimental mutations are, compared to beneficial mutations (Figure 4g), providing a direct comparison with equation (5). We can see a spike in the number mutations that lead to no change in fitness of the whole organism (Figure 4g), which are all the neutral mutations (0.36%), but there are many more detrimental mutations (92.7%) than beneficial mutations (6.94%).

**Figure 4:**
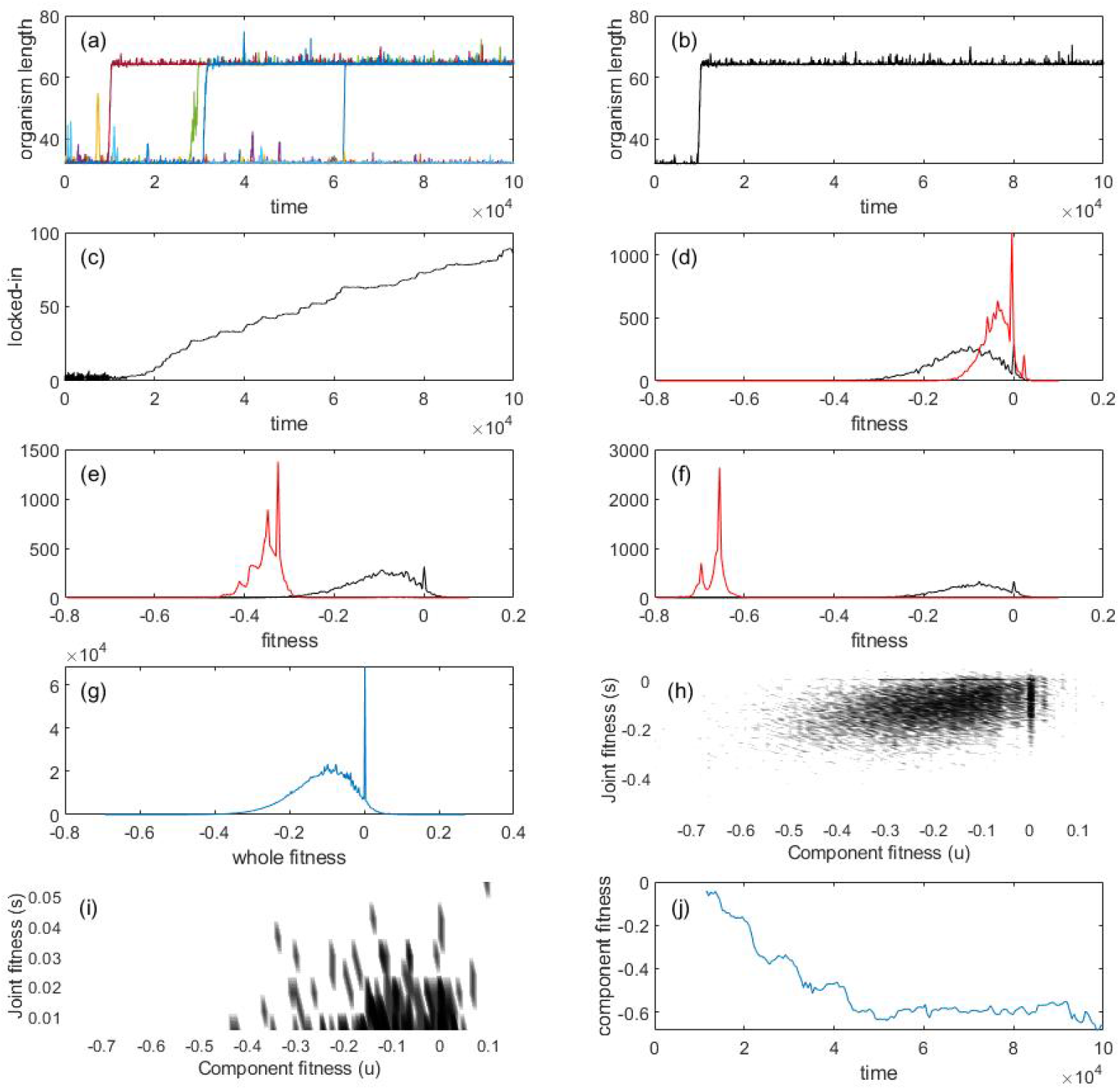
Results from NK simulations. (a) Time series of organism length for eight simulations. (b-j) Results from a single simulation where locking-in of a joining mutation happens early on: (c) Number of links between the first and second halves of the joint organism. (d-f) fitness of all the mutants of the component organisms in the population (red), and of the joint organisms (black) in the population, 100 (d), 1000 (e), and 5000 (f) time steps after the joining mutation. (g) Distribution of a sample of all the mutations that were ever introduced in the simulation (see Appendix B). (h) shows the entire ζ distribution immediately after the joining event. The *x*-axis is the fitness of the component organism, the *y*-axis is the fitness of the whole organisms. Mutants are plotted onto the graph according to the their fitness effect on the whole organism and the components. The evolutionarily important mutations – those that increase the fitness of the whole organism – are plotted in (i; subset of (h) where *y >* 0).

In this experiment, we can also directly measure ζ(*s, u*) right after the joining event and directly see that there is a large class of mutations that benefit the larger agent, while being detrimental to the component agents (Figure 4h,i). This shows the mutation bias that drives the enmeshment: most mutations that are beneficial to the whole organism are detrimental to the component agent. As these mutations become fixed, the fitness of the component agents, *u*, steadily declines over time (Figure 4j). These results are compatible with our earlier prediction that due to the strength of the mutation bias, which may be over an order of magnitude, no naturally observed level of selection is likely to reverse this decline in fitness of the component organisms.

### Generalized evolutionary lock-in

We now generalize this mechanism to a theory of evolutionary lock-in beyond the specific case of joining mutations. In the above example, joining mutations lock in because they create a phenotypic space that was not previously possible – the joint organisms. If this new space is large enough, then supply-driven evolution will push the trajectory of this lineage into the new space, since there will be many more mutations that link the two organisms together, than mutations that separate the two apart. Once in this new phenotypic space (the joint organism), return to the old space is no longer evolutionarily viable I find ‘evolutionarily viable’ a bit vague.

We now generalize this example by defining *σ* as a “potentiation” structure, after Blount et al. (2012) that creates new phenotypic possibilities. To be specific, *σ* corresponds to *N* new mutations, and it is lost *σ* is lost if these *N* mutations become impossible again. We assume this mutation space be unstructured, meaning that mutations in *N* are independent from each other. *σ* becomes locked in when mutations that cause *σ* to be lost become likely detrimental, such that a very long time must pass before *σ* is lost through a beneficial sweeping mutation. If this time is long compared to the duration of the lineage; then the lineage is more likely to become extinct than to lose *σ*. We can compute the expected lifespan of *σ* by assuming that it is proportional to the frequency of *N* mutations in the new space. This is because losing *σ* creates a disturbance in the organism that is itself proportional to the number of mutations in *N*, which reduces the probability that losing *σ* is beneficial.

We now show that *σ*’s lifespan scales exponentially in *N* (to the order of Ω(*a*^*N*^), where *a* is a constant larger than 1), so that even for moderately large values of *N, σ* can lock in. Immediately after *σ* is gained, we first track the number of new mutations *n*, made possible by *σ* that vary from initially 0 to a maximum of *N* . At each selective sweep, the organism can either gain or lose such a mutation, so *n* can either increase or decrease. We do not track the selective sweeps that neither produce a gain nor a loss of a mutation in this space, because such selective sweeps do not change the ultimate outcome in this system. In addition, there is a certain number of mutations, *B*, that causes *σ* to be lost. If *σ* is lost, all *n* of the newly realized mutations are also lost. In the previous NK model, there is only one such mutation (so, *B* = 1), the cleaving mutation that split the joint organism back into two, and all the interactions between the two are lost.

Our task to to show that, for these *B* mutations that cause *σ* to be lost, the probability that they sweep the population becomes vanishing small over time. Let us say that, if *n* = 0 (that is, no new mutations were lost), then the class of all mutations that cause the loss of *σ* has benefit *β*, where benefit is defined as in equation (3): the probability that this class of mutations have of sweeping (both being advantageous and escaping drift). In the NK model, this would be the likelihood that a cleaving mutation sweeps the population if there were no links between the first and second halves of the joint organism.

We re-emphasize that, when *σ* is lost, all the mutations that were gained in the new mutation space are lost. The more mutations that are lost, the less likely that the loss of *σ* would be beneficial. If *n* new mutations were lost, we assume that the probability that the reversal mutation is advantageous is *μ*^*n*^ for some 0 *< μ <* 1. We justify this as follows: let’s say that for the *n* mutations lost, each lost mutation has a *ν* probability of being advantageous. Conservatively, let’s say that more than *n/*2 of the mutations lost must be advantageous for the mutation that reverses *σ* to be, overall, advantageous maybe conservative but also arbitrary. How sensitive your results would be to this assumption?. This corresponds to the cumulative binomial density function, Pr(*B*(*n*, 1 − *ν*) ≥ *Ln/*2*J*). Since *ν* is the probability of an advantageous mutation, we can reasonably assume that 1 − *ν >* 1*/*2, which means that Hoeffding’s inequality holds you should provide some context for this inequality:

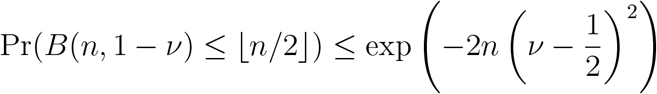

The probability that a mutation that reverses *σ* is beneficial is at most *μ*^*n*^, where

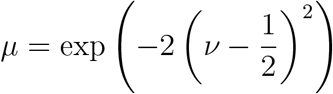

Thus, the probability that a mutant which lost *σ* can sweep the population is simply *βμ*^*n*^.

At time *t*, let there be *n*(*t*) new mutations that have been actualized. At this point, there are *n* new mutations that can be lost and *N* − *n* new mutations that can be gained. If all mutations have the same probability, this corresponds to the force this is too obscure: what is ‘this’ and can you be more specific about the ‘force’? of supply driven mutation that will increase or decrease *n*(*t*).

There can, of course, be an environmental side to all this, such that increases or decreases in *n*(*t*) can be favored or disfavored by the environment. However, this will be unimportant in our case for even moderately large values of *N*, since at the beginning of the process of exploring the new mutation space, *n* will be very small and so *N* ≫ *n*, a difference that will be very hard for the environment to overcome. By the intuition we have developed in this paper, we can see that although the environment can alter the specific values of equilibrium *n*(*t*), it is unlikely to alter the ultimate outcome of whether *σ* can be locked-in.

In this case, let us assume that the fitness benefit of increasing *n* to be *p*, and of decreasing *n* to be *q*. The benefit of a mutation that reverses *σ* is *β*. With these notations, we can now model the system with the Markov chain transition matrix, **S**, of size *N* + 2:

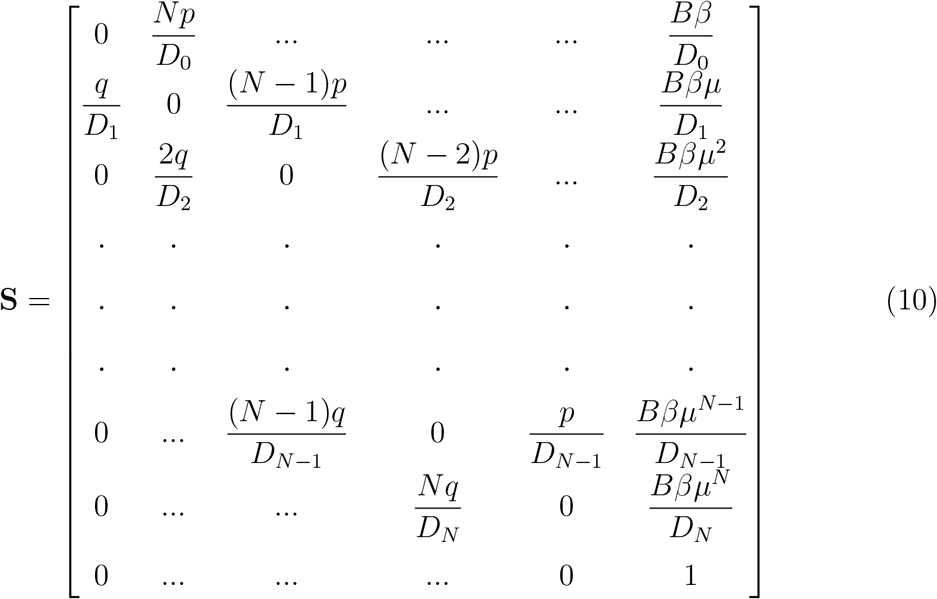

Where *D*_*i*_ = (*N* − *i*)*p* + *iq* + *Bβμ*^*i*^.

We can show that the expected time to losing *σ*, or *T*_*B*_, for *p* = *q* = *β* and *B* = 1, is (see Appendix C)

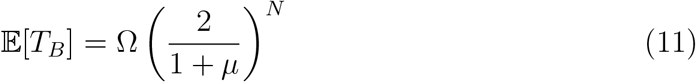

This does not qualitatively change if we alter the values of *p, q* or *β*, for any constant *B*, once a potentiation structure comes about, the expected time before it is lost increases exponentially with its amount of phenotypic novelty. We should also note that this process does not only involve the opening of new phenotypic spaces by the introduction of *σ*, it also closes off old phenotypic spaces, because all the mutations that cause *σ* to be lost (there are *B* of them) are now rendered detrimental. There is thus both an opening, but also a closing of evolutionary space over time. there is here some ambiguity about mutations that are lost (no longer exist) and mutations that are detrimental. This mechanism works generally: all evolutionary innovations that opens sufficiently large new phenotypic space can become locked-in. *σ* can represent the formation of eukaryote, multicellularism or eusociality. I can also be associated with new ways of patterning that are not necessarily hierarchical, such as the formation of body axes, or or limbs.

## Discussion

For a long time now, some evolutionary biologists have had an intuition that macro-evolution is a different kind of process from micro-evolution. The mechanism responsible for this difference have been debated: they range from species-selection (Vrba, 1984; Jablonski, 2008), punctuated equilibrium (Gould, 2004; Eldredge et al., 2005), to processes that only happens when there is no natural selection (the “zero-force evolutionary law”) (McShea and Brandon, 2010), and others. All of these mechanisms have been criticized from the view that macro-evolution is “nothing but” successive rounds of micro-evolution (that is, selection and drift), with some accommodations made for higher hierarchical versions of micro-evolutionary process, without genuine theoretical novelty.

Balanced against the latter view is a nagging suspicion that there are phenomena in macro-evolution that are nothing like micro-evolution at all; issues of historical contingency such as Gould and other paleontologists articulated, calling for evolutionary biology to be an essentially historical science (Gould, 1989; Beatty, 1995, 2008), events such as the major transitions of evolution (Szathmary and Smith, 2000; Szathmáry et al., 2011), and troubling issues of evolutionary lock-in where events early on in time are “locked-in” for the rest of that lineage – there are no six legged tetrapods (Schank and Wimsatt, 1986; Wimsatt, 1986; Klingenberg, 2005). These problems are stubbornly resistant against standard explanations of micro-evolution. In this paper, we have shown a biologically and mathematically plausible mechanism that might provide an answer to these problems. the following statement is central and needs to be more specific This view reconciles macro-evolution to microevolutionary processes, such that macro-evolution is driven by micro-evolution (natural selection), but over many episodes of micro-evolution, new patterns and trends appear, driven by the way new evolutionary spaces open and close.

To do so, we first defined an evolutionary regime called “supply-driven evolution” (SDE), where mutation bias is the chief driver of the evolution of certain traits.

The first requirement for SDE is that the trait in question must be realizable in many different phenotypes. For example, two very different organisms can have the same height, weight, or complexity however measured. Secondly, we assume evolutionary change comes about primarily by the selective sweep of rare beneficial mutations. When these two conditions are satisfied, we showed that mutation bias becomes as important as natural selection, because the order of introduction of mutants can now determine the direction of evolution. This is because if a particular trait value is introduced more frequently, then it is more likely to succeed.

Once these conditions are satisfied, we proposed that there are scenarios where mutation bias can be so strong that no environment is likely to reverse it. We called these regimes “supply-driven” evolution (SDE). One such scenario of SDE can drive a number of the major evolutionary transitions: the steady and seemingly difficult to reverse increase in structural hierarchies, from self-replicating RNA molecules to prokaryotes, to eukaryotes, to multicellularity, to eusociality. To explain this, we defined as our trait the likelihood of breaking a joint organism into its components, such that the components are more fit than the joint organism. This trait is clearly realizable in many different ways, and we showed that there is a strong mutation bias that drives this likelihood down over time. This is because there are more detrimental mutations than beneficial ones, such that most of the mutations that benefit the joint organism will be detrimental to the component organisms.

We then showed that this mechanism generalizes to any structure that makes possible new phenotypes. If a new structure came about that makes available a new set of possible phenotypes that were not previously evolutionarily accessible, then we showed that this structure can be evolutionarily locked-in over time such that no descendent of the lineage of this organism will break it. The measure we consider is the likelihood that this structure will break in a way that is selectively favorable; but there is a strong mutation bias driving this likelihood lower over time. The bias is because there are, at least initially, many more ways to enter into the new set of possibilities than to leave it, and once enough new possibilities is realized, breaking the structure that made it all possible is almost certainly detrimental.

A large number of macro-evolutionary phenomena can be addressed by this theory. The hox-genes, for example, by allowing for head-tail patterning, might have created a host of new evolutionary possibilities that were not possible before. As those new possibilities became realized, hox-genes could not longer be replaced. Similarly for the hemoglobin protein: once the most evolutionary optimal (or at least locally so) protein carrier of oxygen was found, it made possible many new possibilities that were not available to lineages of lower metabolism. All the structures made possible by increased metabolism would be broken if the hemoglobin became less efficient. It is also easy to imagine that the capacity for human culture is itself evolutionarily locked-in by the realization of new possibility. Although many objections could be made to these just-so explanations, the least of which is that there is little evidence (something we will address below) the theory of SDE nevertheless expands our theoretical imagination beyond the Modern Synthesis, while not rejecting any element in the Synthesis. The fact that eukaryotism has persisted might have nothing to do whether it is evolutionarily advantageous for the lineage (except for that short moment in which it came about); but rather because eukaryotism was impossible before and possible afterwards. Once a large enough swathe of novelty becomes *possible*, SDE shows that there is a chance the structure behind the novelty becomes necessary.

Another important phenomenon can be made sense of in the light of this theory: the empirical insight that the most evolutionarily impactful innovations are actually quite silent in the moment of their invention. This was recently reviewed in Erwin (2021), and they sought to explain this phenomenon in a manner very similar to the one given here:

Novelties generate the design or opportunity spaces in which subsequent adaptive radiations may occur, but the origin of novelties is largely decoupled from subsequent evolutionary diversifications.

In Erwin (2021), however, they did not give a theoretical justification why there is this initial silence, or why such innovation persists for a long period of time. We formalize this intuition and show how it is entirely compatible with microevolutionary theories of selection.

In other words, the macroevolutionary impact of an evolutionary change is not the ecological effect it has in the moment, but the size of the novel evolutionary space the lineage gains. A large space “sucks” the lineage into it, and its future ecological effects take place inside the new set of possibilities. In this way, the opening of new evolutionary space becomes the macro-evolutionary equivalent of fitness; a new structure that opens new evolutionary space has a probability of “locking-in”, in the same way that a mutation that increases fitness has a probability of sweeping. This is happening even when all that observers can see are selective sweeps of populations through differences in fitness. What the observer cannot observe, however, are the enormous spaces of mere potential that might be opening or closing at each selective sweep. On the other hand, these invisible spaces exert a real evolutionary force, metaphorically similar to a vacuum, because a large potential space will “suck in” evolutionary trajectories. Such a large space form the underlying and invisible universe of the possible from which the actual and observed are chosen.

This is a macro-evolutionary process since the process takes place over many successive selective sweeps, and the progression of the lock-in is more or less uninterested in environmental change.

This way, evolutionary biology might become a truly historical science. Depending on what order of mutations came along, vast spaces of new possibilities might be created while potentially other, even larger avenues of historical possibility are simultaneously destroyed for all futures of that lineage. We suspect, but have not shown, that *more* historical possibility can be destroyed than the new possibilities that come into place, which can create a phenomenon of evolutionary stasis.

Other examples of this historicity further stretch our imagination of what is possible. For example, it is well known that most mammalian lineages have seven vertebrae (Galis, 1999) such that alterations of this number is significantly teratogenic – the biological consequence of having too much broken all at once (Galis et al., 2006). Only mammals with low metabolism, such as the sloth and the manatee, have escaped this restraint. Thus, having seven cervical vertebrae was somehow necessary to the development of mammalian metabolism. This shows us that the mechanisms by which new evolutionary possibilities are created, and what potentiation structures can get locked in over time, are not at all obvious.

The theory advanced by this paper has had many predecessors. Besides the direct influence of the authors cited in the Introduction, there is much that was presaged by Gray et al. (2010) (constructive neutral evolution) and similar work (e.g. Force et al. (1999)): complex structures can be created by duplication, each duplicate of which accumulate deleterious mutations. This is clearly similar to what this paper propose happens in the evolution of hierarchies: components join together and can longer depart because of the accumulation of deleterious mutations. Parts of the work here was also inspired by McShea and Brandon (2010), where McShea clearly noticed that there are more ways to be different than to be the same – and that this can form a force to drive organisms to become more complex. Finally, the idea of a topology where evolutionary possibilities sit on a thin web linked by threads was advanced by Crutchfield and Van Nimwegen (2002) as well as Gavrilets (1997). However, this paper is different from all these previous work because it makes the insight that such phenomena are compatible with regimes of high natural selection, which can actually advance these dynamics. SDE is thus able to roots its micro-evolutionary process on selective sweeps, while all other works are based on neutral evolution. Supply-driven evolution works regardless of the strength of selection; in fact, the stronger the selective regime, the stronger the theory proposed by this paper, because at any one point there are fewer evolutionarily relevant mutants, making the order in which mutations come about become increasingly important.

From a wider theoretical vantage point, there are two other close predecessors. One comes from Kauffman (Kauffman, 1993), in what he called “ensemble theories” of biology. For example, he remarked that the number of cell types that any organism had might be eerily similar to what a “typical” organism with that number of genes would be expected to have. Without going into the details of his model, we can see how this ensemble style of argument can work with SDE: among all possible types organisms with a certain number of genes, the largest “ensemble” will have, by definition, the typical number of cell types. The insight that SDE makes is that that the fittest organism is also most likely to lie in that ensemble. Kauffman hypothesized that natural selection is unlikely to change the cell number; we can now justify the hypothesis with the insight that natural selection might be very strong, but it is acting for or against specific forms of the organisms – and the fittest form is mostly likely to lie in the largest ensemble, that of the “typical” cell type number.

The second predecessor is Gould’s loosely argued insight that structure is integral to biology, but is left out of the Modern Synthesis. This was most forcefully brought up in his last book Gould (2004), but appear through many of his other works as well. SDE captures some of his insights. After all, the innovation that an organism can generate is highly dependent on the existing structure of that organism: what those structure make possible, and make impossible. For example, exaptation has a natural home in SDE; a structure that creates a new space of possibility that depends on itself is also creating new role for itself: for supporting the new space of possibility. Feathers, for example, originally selected for temperature homeostasis, also creates a new space of new possibility, which includes flight and, from there, the entire evolutionary space of birds, which were impossible prior to feathers. SDE shows that when such a new space is created, the innovation locks in and its continued existence becomes divorced from the original reason the innovation was selected for.

### Theoretical directions

This view of evolution raises deep questions. The primary problem is that we have asserted the existence of mutations that make new evolutionary space possible. This seems intuitively plausible for the biologist, but it is not theoretically obvious. To assume it is to assume a certain topology of permissible evolutionary space, one that resembles a tree - such that there are moments where taking a particular direction opens new fields and closes old ones. In wide-ranging papers, Crutchfield (Crutchfield and Van Nimwegen, 2002; Crutchfield, 2003) hypothesized precisely this type of space to drive what they called “epochal evolution” and they demonstrated several fitness functions that satisfy such a topology. However, no further justification was given why this topology would be the one we see in nature.

In essence, by asserting there exist mutations that create new evolutionary space, we are asserting that life is based on an intricate network of dependencies - structures work because they depend on each other. To a biologist, this might seem obvious, but to a theoretician, it is obscure. What sort of systems demonstrate this dependency? What sort of structures create new evolutionary spaces? Are all such structures frozen after structures that depend on them evolve, or can they still be changed, with their own evolutionary trajectories? What is the distribution of the sizes of the new evolutionary spaces that they potentiate? Can they be predicted in advance? All of these are open questions.

### Empirical directions

We freely admit that there is currently not much experimental evidence for SDE, although we would argue that the history of life and what we know of biological processes are consistent with such a theory. Our existing experimental data have been driven by questions asked by the Modern Synthesis and are thus limited in determining the phenomenon of SDE.

There are nevertheless tantalizing clues. In Lenski’s (Blount et al., 2008; Wiser et al., 2013; Blount et al., 2012, 2018) now classic and ongoing Long Term Evolution Experiment (LTEE), they discovered a fascinating new innovation, aerobic citrate utilization among their lineages of *E. coli*. This is so unusual that the inability to do so is considered a defining feature of wild *E. Coli*. Nevertheless, it was discovered in the LTEE, and their team studied why this innovation was so rare. They discovered that, in order for this ability to use citrate to come about, several other mutations had to first be in place – a process they called “potentiation”, which was later “actualized” by the capacity to use citrate.

While the actualization mutations were carefully isolated and studied, the potentiation structures prove much more difficult. As Blount et al. (2012) states:

The potentiating mutations are not known to confer any phenotype amenable to screening, so there is no simple way to distinguish between potentiated and non-potentiated clones…

Despite heroic experimental work, Blount et al. (2012) was only able to determine that potentiation involved at least two mutations, the approximate timing of these mutations, and evidence for the hypothesis that these mutations potentiated the actualizing mutation through epistasis (gene-gene interaction), rather than simply rearranging the genome so the actualizing mutation became more physically likely.

This fascinating work shows that SDE is, at least, theoretically possible. Potentiation structures exist. In that sense, our paper shows that a potentiation structure, if it opens a large enough space, will likely lock-in and form a part of historical contingency (Blount et al., 2018). However, Blount *et al*.’s work also shows the current experimental challenges to SDE: potentiation structures can be evolutionarily very quiet. Whatever the size of the space they have opened up, that space remains silent until the actualization, and observers only see the actualization – the rest of the unactualized space is never observed, even if that unactualized space exert a powerful effect on evolution by changing the underlying potential universe from which new mutations are selected.

To study SDE experimentally, the first step is relatively easy. It would involve constructing a mutation library of as many mutants as possible of a certain lineage, then classifying this mutation library according to the phenotype of each mutant: its fitness and whichever trait we are interested in. Such a library is already possible, for example, it was done for the important recent paper on mutation bias (Monroe et al., 2022), as well as older work such as on the tomato plant (Menda et al., 2004). From this library, we would then construct the fitness distribution of these mutants, as plotted against the trait value, to construct ζ. One can then subject this lineage to evolution, and see if the trajectory of the trait in question follows the prediction of the framework laid out in this paper.

Importantly, if such a library is constructed at multiple moments of the evolution of this lineage, it would establish whether ζ stays stable over time - an open empirical question. Such an mutation library can at least answer the question of whether long term trends in evolution through mutation bias is possible.

The next step, the search for potentiation structures that open up evolutionary space, is trickier. Just like above, we would have to be able to construct successive construction of mutation libraries, which are approximations of the underlying potential universe from which new innovations are selected. However, more than the above, we would have to be able to sequence and understand the changes in structure of the mutants in those libraries, such that we can identify changes in these mutation libraries. We would look for instances where previously impossible phenotypes are now possible, or previously accessible phenotypes have now become lethal or cannot be generated. This would provide a view of the how frequent potentiation structures are, how quickly the underlying universe of possibilities change, and how likely is evolutionary locking in over time.

### Conclusion

We hope that SDE can significantly shift the discourse on highly contentious topics of evolutionary biology, such as the evolution of altruism, culture, intelligence, and other traits of human interest – or that make humans special. Instead of taking a selectionist view that carefully study the environmental conditions in which certain structures (or behaviors) are promoted and maintained, we might instead want to ask what how new mutations are generated, and what new possibilities are created. Such a view of evolution, and indeed, of culture, would give novelty and innovation their proper place in evolutionary theory. Perhaps evolution is biased not only in favor of survival, but also in favor of creativity.

Beyond that, we can gain a scientific understanding of historical contingency, instead of relegating this vast phenomenon under the rubric of “chance”, or simple path-dependency. There is the hope of finally understanding two classes of change, one that is potentially historically significant, and one that is not. Potentially historically significant change might be those where new structures come about that open up large spaces of new possibility. These are the structures that have a chance to be frozen in time – not to be changed for all descendants of a lineage. Historically insignificant structures do not open up new space. However useful they are for the moment, insignificant structures can be discarded when the environment changes. These and other possibilities would certainly enrichen modern evolutionary theory.

## Appendix A: Details of the NK model for supply-driven evolution

We define an NK model that is similar to, but different from, the model described in Xue et al. (2015). Agents are now strings of *L* bits, either 0 or 1. Each bit is an “element”. Each element is linked to *K* other element in the agent, forming *L* substrings of *K* + 1 length. Before running the model, a “fitness table” is generated. In this table, a fitness value is associated with each possible substrings of *K* + 1 length. The agent’s fitness is simply the average fitness of all its *L* substrings, by looking up each substring in the fitness table. Since fitness in this landscape depends on elements that are linked together, the effect of any one component value depends on other element values. In the original NK model, the fitness table is populated by fitnesses that are randomly chosen from 0 to 1.

Unless stated otherwise, the population consists of 5000 agents, each of which is a string of *N* = 40 elements. *K* is set to 3. The reason for this is is detailed in the next paragraph. All agents are wired identically in a wiring diagram that is chosen at the beginning of each model run. Each element has equal chance of being connected to any other element in any order. Element values is either 0 or 1. The fitness table represents the environment and will be specified below. When the environment changes, this change is represented by choosing a number of entries from the fitness table and changing the fitness values associated with these substrings. The rate of environmental change is the same for all models, at each timestep, each entry in the fitness table has 10^−5^ probability of being changed.

The reason *K* is set to 3 is because if *K* is very small, the fitness table is also very small (it is, after all, simply 2^*K*+1^). For *K* = 2, the fitness table is only 8 in size. This means that any environmental change is very large in scope, as the fitness of 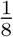 of all substrings are changed. This introduces a wild source of noise into our runs, which can be seriously altered with any change to the fitness table. With *K* = 3, this noise is much reduced. Higher levels of *K* slow down the run. For all values of *K >* 1, however, the outcomes are qualitatively the same.

For the run itself, at each timestep, *N* agents are chosen with replacement for update. An agent is selected with probability proportional to its fitness to reproduce one other agent. Another agent is then randomly selected to die. This is a Moran process. Thus, we can set *α* everywhere in our equations to be equal to 1. The descendent is identical to agent *k*, except for mutations. Each element of the parent agent *k* has 10^−5^ probability of being different in the child. If a element mutates between parent and child, the value of that element will either increase by flipping from 0 to 1, staying as 1 if it already was 1, or decrease by flipping from 1 to 0, staying as 0 if it already was 0.

We will study the evolution of the trait 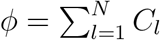. We generate a fitness table such that phenotypes with higher trait values have, on average, higher fitnesses. For any substring *s*_1_*s*_2_ … *s*_*K*+1_, its entry in the fitness table is a random number chosen from the exponential distribution with the mean 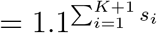. Thus, each increment of 1 in the substring leads to a 10% increase in fitness. Overall, then, an agent with larger values in each of its elements will also have higher fitness. However, the variable fitness assumption is satisfied because many different agents can have the same trait value, but have different fitnesses.

A mutation bias is here introduced where there is 90% chance that, if a mutation happens, the element decreases. There is thus a 10% chance that, if a mutation happens, the element increases.

Each time an entry from the fitness table changes due to environmental change, it is re-sampled from the same distribution, *λe*^−*λx*^. Here *x* is not the same as *ω* as in the previous models, since *x* denotes only the fitness contribution of a substring to the whole organism’s fitness, not *ω*, which is the organism’s whole fitness. This does not happen if there is no environmental change.

This specifies the NK model used in the paper.

We ran this model with the above specified parameters for 200 runs, for 1e5 time steps each. We ran this model for another 200 runs without mutation bias. Final results can be seen Figure 3.

We also present Figure 5, a figure for individual results of each of 200 runs with mutation bias.

**Figure 5:**
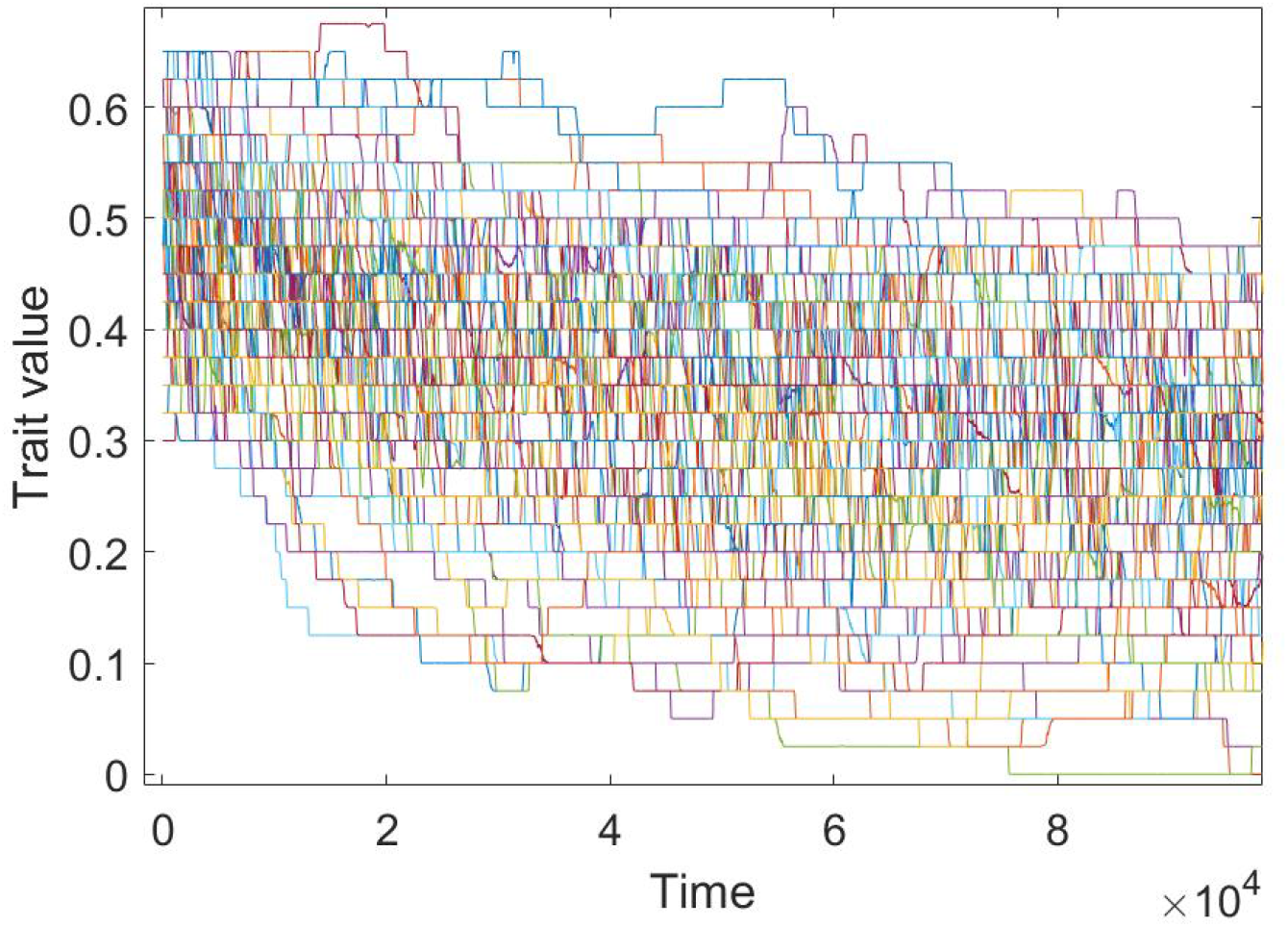
All 200 runs of the model

An interesting side effect of mutation pressure depressing the trait value over time is that the average population fitness over time can fluctuate and have long periods of decrease over time (Figure 6).

**Figure 6:**
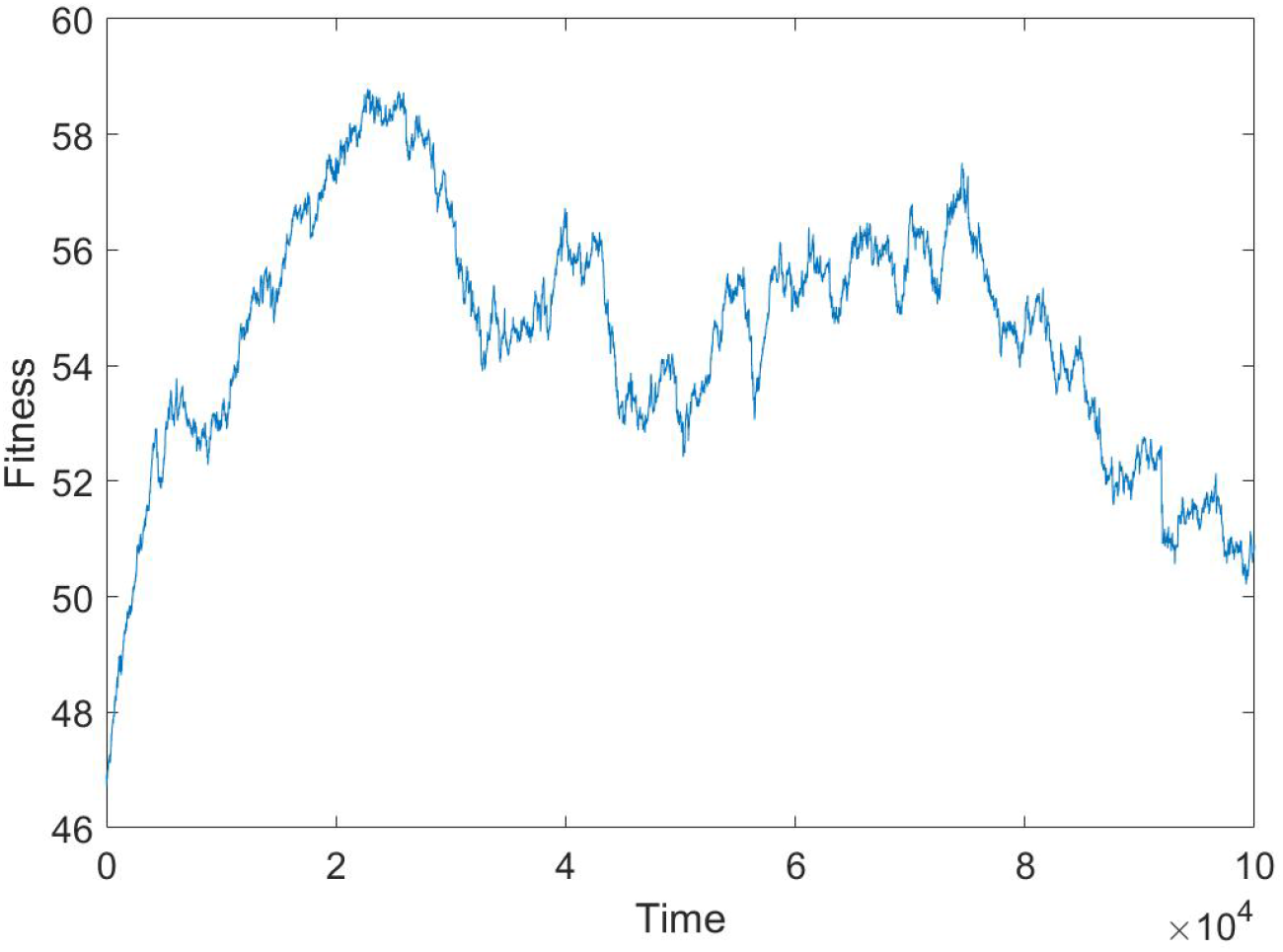
The mean fitness change over all 200 runs of the model

**Figure 7:**
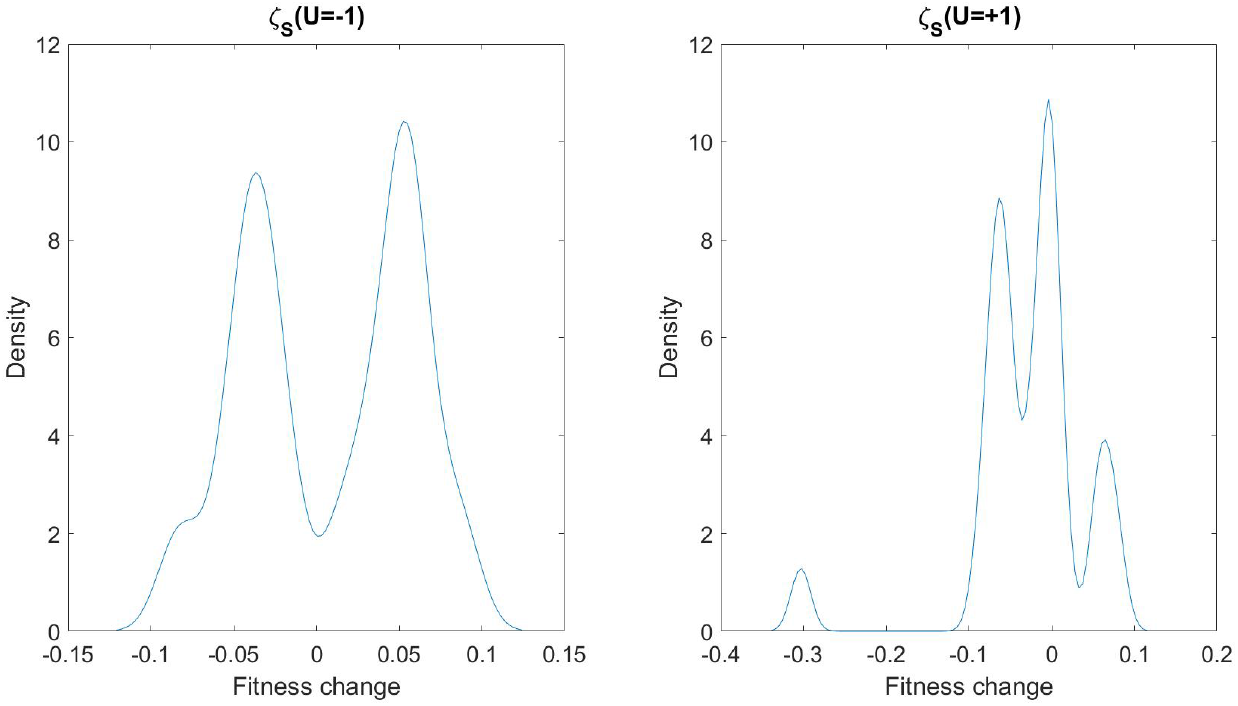
An example of the ζ_*S*_ distribution of one of the model runs, at t=1000. The panel on the left is the fitness distribution of all the mutants that decrease trait value by 1, the panel to the right is the fitness distribution of all the mutants that increase trait value by 1.

To test our theoretical predictions, let *ϕ* be our trait value, which is the number of 1’s in an organism, while *N* − *ϕ* is the number of 0’s in an organism. We observe the following: excepts for rare spurts of selective sweeps, the population usually has a dominant, homogeneous wildtype (see Figure 5). The *ϕ* value of the wild type is more or less the population *ϕ* value. In this population, then, ζ_*U*_ is simply made from two parts: probability to 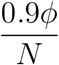 increase the element value, and probability 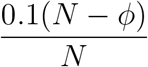 to decrease the element value. As the population evolves and *ϕ* changes, ζ_*U*_ also changes.

The distribution of benefit *β*(*u*) at any one point can be simply obtained. At pre-defined times (in the runs, they are measured 10 times over the entire run at equal intervals), 10000 mutants of the wildtype are examined, without mutation bias – 5000 of these mutants decrease trait value and 5000 of them increase trait value (mutants that does not change trait value are not examined). These mutants are not real in that they are not a part of the population, they are only used for calculating *β*. The fitness of each of these mutants are noted and ζ_*S*_(*s*|*U* = *u*) is constructed. An example of a ζ_*S*_ is shown below, for one of the runs at time = 10000. Since there are only two possibilities for trait value change in our model, we present the two distributions ζ_*S*_(*s*|*U* = −1) and ζ_*S*_(*s*|*U* = +1):

From this ζ_*S*_, it is simple to compute

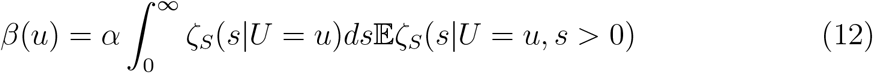

per our definition of *β*, using *α* = 1 since we implemented a Moran process.

By computing *β* is this way at a certain time step, we can attempt to predict how the trait value will evolve, since we know how ζ_*U*_ changes. We assume *β* remain relatively constant. We can then use these parameters to predict the evolution of the trait value *ϕ* in the model. We choose *t* = 10000 as the time to measure *β*, as that is the moment where the population had settled down to some sort of equilibrium with the environmental change. The results are seen also in Figure 3. The fit between theory and agent-based model is good rather than perfect, showing that *β* does change over time, but better work in predicting how it changes over time will have to be in the future.

## Appendix B: Details of the NK model for the rise in hierarchies

We will detail here the NK model used to show how increases in hierarchical organization can be irreversible.

We start with a population similar NK model as in Appendix A, with a population of 2000 agents, evolving in a Moran process. The initial *N* for all these agents, however, is 32 (we use exponentials of 2 for *N* so they can be readily cleaved or doubled). The initial *K* is much larger than before, *K* = 10, we will see the reason for this below. The environment still changes as above, there is a probability of 10^−5^ of any entry in the fitness table being changed at any timestep. The fitness table is now a lot larger, of course, due to *K* being so large.

Mutations in the previous model changed the element value of the agent from 0 to 1 or vice versa. Here, there are no mutations that change element value - whether an element is 0 or 1 is set at the very beginning of the run with 50% chance of each. Instead, mutations in this model change the wiring between two elements. As we have seen, a NK model is defined by its elements as well as the wiring among those elements. Thus, a mutation in our model will take a link between two element *i* and *j*, and throw the second element to a random other element, *k* ≠ *j*. This potentially changes a substring in the organism and thus potentially changes its fitness.

There is an issue with implementing mutation in this straight forward way, however, because we will see that the number of links in this model can change over time through joining and cleaving mutations (detailed below). Thus, the effect size of a mutation changes based on the number of wiring that the agent has. In order to normalize this effect, we define a probability, *probRewire*, in our model, such that when an agent is to be mutated, all of its links have this probability to be rewired. We set *probRewire* = 0.005 for our model runs. This means approximately 2.5% of all the substrings in an agent is changed whenever the agent is mutated.

There are two other types of mutation that happens in this model, joining and cleaving mutations. The probabilities of these two types of mutations are set at 0.025 each, such that 0.95 of all mutations changing the wiring of an agent. Thus, when a mutation happens, there is 0.025 probability that two organisms join together and form a single joint organism, and the same probability that an organism is cleaved into its halves, the two component organisms. When a cleaving mutation happens, all wiring between the two components are lost. In order to keep *K* the same, these wiring are thrown to random other elements in the component organism.

The three types of mutations are mutually exclusive. In order to keep the population size the same, when there is a cleaving mutation, one of the components is randomly removed, and when there is a joining mutation, one of the two original organisms is randomly retained.

Because of the way that the cleaving mutation is implemented, its effects are mitigated

There is one other issue of implementing joining and cleaving mutations in the straightforward way. The problem is that, given an existing wild type, there is exactly one possible outcome of a joining mutation, and exactly one possible outcome of a cleaving mutation. This is a violation of the multiple fitness assumption, that there are many different ways of joining or cleaving in length, each with a different outcome in fitness. This means that there is no mutation bias effect possible in the joining or cleaving events, since at any one point, the fitness outcome of a joining or a cleaving event is predetermined. It also makes several of the results below difficult to analyze. To overcome this, we added a process whereby after any joining mutation or cleaving mutation, there is also a rewiring mutation, where 0.5% of all wires of the joint or component organisms are mutated. This effectively means that there are many ways for joining mutations to happen, and many different ways for cleaving mutations to happen. This probably corresponds to biological reality: there is more than one way for increased hierarchies to evolve, and there is also more than one way for hierarchies to dissolve.

This completes our description of this model. To generate the results we see in Figure 4, we run this model 8 times for 100,000 time steps. For panels 4d-f, we took one of the runs in which joining happened and was locked in place, and we measured the fitness distribution of all the mutants of the joint organism (black line), as well as the fitnesses of the component organisms that would result from a cleaving mutation (red line). We do so by examining 10,000 mutants of the joint organism (wiring only), and 10,000 component organisms that results from a cleaving mutation after the wiring mutations. We do so three times: 100 time steps after the joining mutation manages to sweep the population (Figure 4d), 1000 time steps after the joining mutation has swept (Figure 4e), and 10000 time steps after the joining mutation has swept (Figure 4f). We see that the red distribution becomes less and less fit as time goes on. We see that after sufficient time, the fitness of the component organisms decay so much that it becomes basically impossible for any component organism to sweep the population. The decay of the average fitness of the component organism comes can be seen in Figure 4j.

Figure 4g-i show the mutation bias that drive the system. Figure 4g show the fitness distribution of 10,000 mutants of a joint organism (wiring only). Many of the mutations are neutral, but the vast majority are detrimental and less than 7% are beneficial. Figures 4h and 4i show the empirical ζ in the system. We examined 10,000 mutants (wiring only) of the joint organism and their effects on the fitness of the wild-type (y-axis), as well as their effect on the component organisms (x-axis). Figures 4h shows the overall ζ distribution, Figure 4i shows only those mutants that are advantageous for the joint organism. We can see that most of these mutations are disadvantageous for the component mutant, which means that the fitnesses of the component organisms decrease over time.

## Appendix C: Evolutionary locking-in by the discovery of new evolutionary space

We start with equation (10). For simplicity’s sake, we first give the proof of the case where *p* = *q* = *β* and *B* = 1. This gives the basic methods of the proof that can be generalized later. In that case, equation (10) becomes:

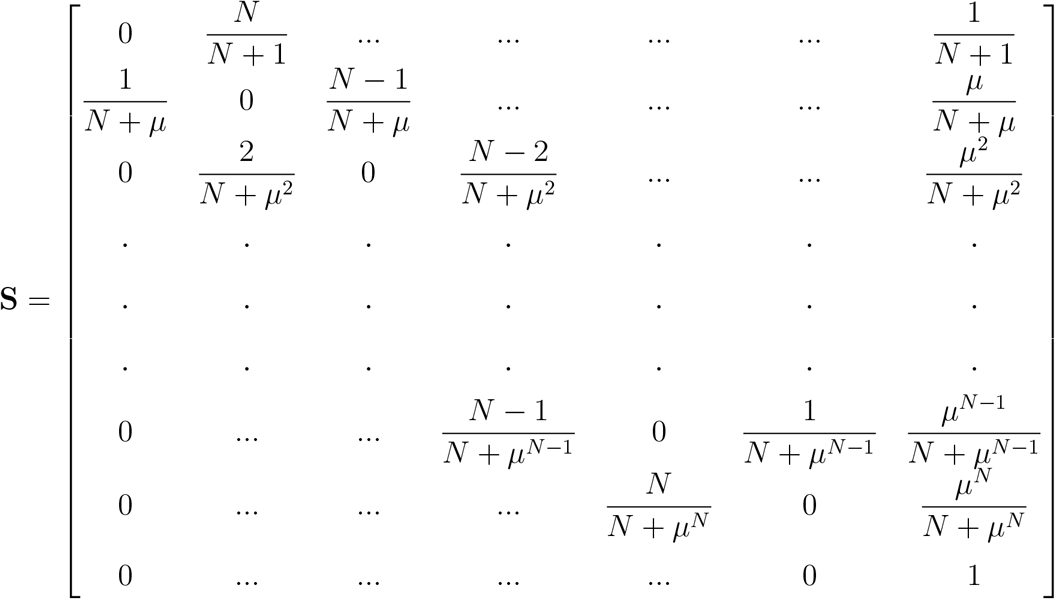

In **S**, all … represent zeros. The first *N* + 1 rows represent organisms with *n* new mutations (0 ≤ *n* ≤ *N*) that are realized. We discount mutations that simply return *n* to *n*, hence the zeroes on the diagonals. The probability of increasing *n* is simply 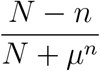, of decreasing *n* is 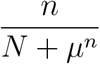, and of successfully (that is, selectively advantageous) breaking *σ* and destroying all *n* mutations is 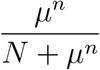. *β* = *p* = *q* cancels out through the numerators and denominators. In reality, *β* becomes a time scaling factor – it measures the time between two selective sweeps, so that time flows faster for larger *β* and slower for smaller *β*.

The last row in **S** is the absorbing state. If *σ* is destroyed, we make the conservative assumption that it stays destroyed and cannot come about again. This makes our system a discrete phase-type distribution and is entirely characterized by **S** minus its last row and last column, knowing that all the rows sum up to 1 and the last row is the absorbing state:

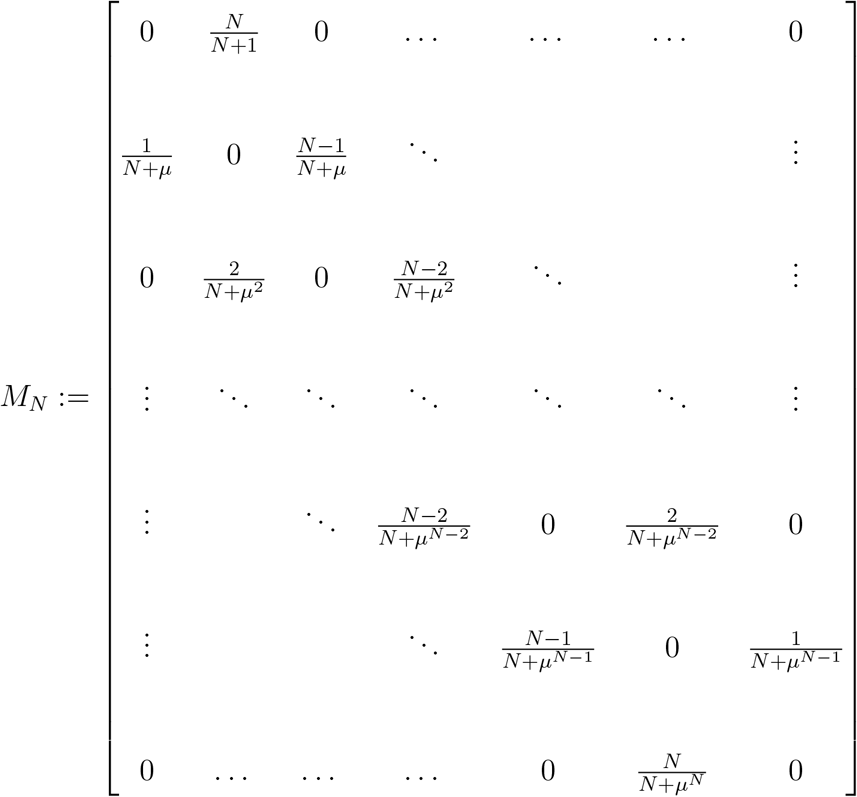

and its starting state is the vector *α* = [1, 0, …, 0]. By the standard theory of absorbing Markov chains, the quantity of interest, the time until the system reaches the absorbing state, is given by

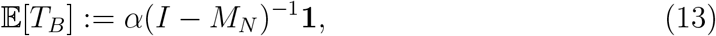

where *I* is the identity matrix and **1** is the vector of all ones.

### The upper limit case: μ = 1

The simplest case of (13) arises in the limiting case *μ* = 1. When this occurs, the matrix *M*_*N*_ can be written as 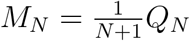, with *Q*_*N*_ **1** = *N* **1**. Hence, the equation

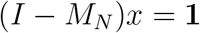

has the obvious solution *x* = (*N* + 1)**1**, meaning that 𝔼[*T*_*B*_] = *N* + 1 (in fact, this is true no matter what the initial state is). This means that, when the probability of breaking *σ* is a constant independent of the number of mutations *n*, the expected time to reversal is proportional to the number of possible mutations.

### 0.1. The lower limit case: μ = 0

In the second simplest case of (13), when *μ* = 0, we also have identical denominators throughout *M*_*N*_, except for the first row:

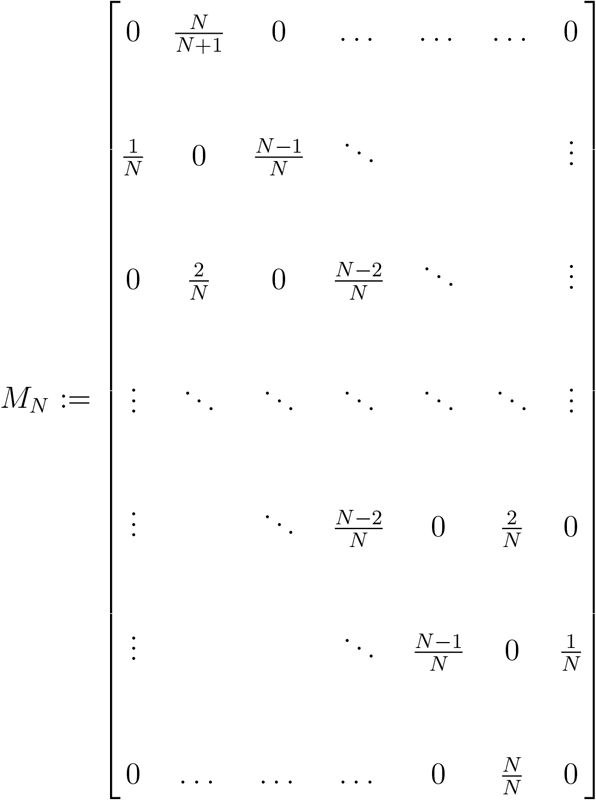

The strategy we will adopt is to solve the system (*I* − *M*_*N*_)*x* = **1** from the bottom up. Numbering the entries of *x* from 0 to *N*, the last equation in the system simplifies to

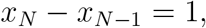

the first one simplifies to

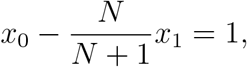

and the remaining ones take the form

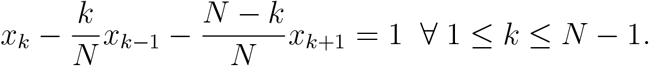

While solving the system in decreasing order of *k* values, it is easy to see that in fact, the solution satisfies

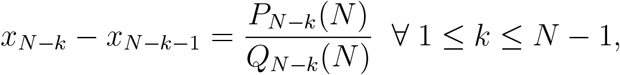

where both *P*_*N*−*k*_ and *Q*_*N*−*k*_ are polynomials of the same degree *k*. In fact, the explicit form for *P*_*N*−*k*_ and *Q*_*N*−*k*_ is given by

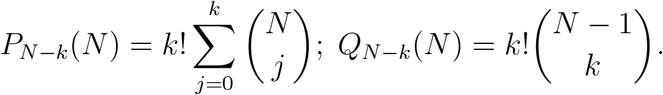

We now show both of these statements simultaneously by induction on *k*. The base case, *k* = 0, results in *P*_*N*_ (*N*) = 1 = *Q*_*N*_ (*N*), which is consistent with the initial equation *x*_*N*_ − *x*_*N*−1_ = 1. Suppose that this result holds for some value *k* − 1 with *k >* 0. Substituting it into the (*N* − *k*)-th equation in the system yields

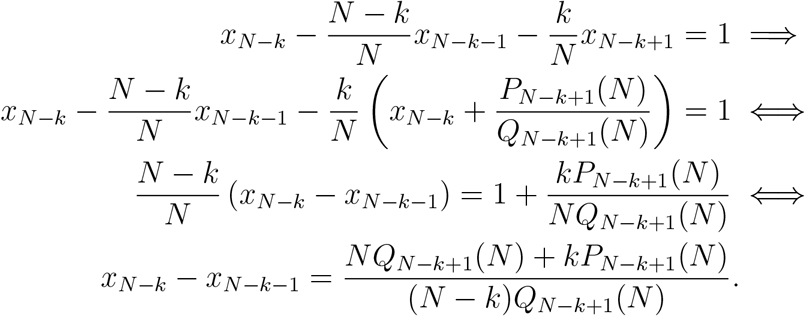

The desired statement now follows from observing that the denominator is

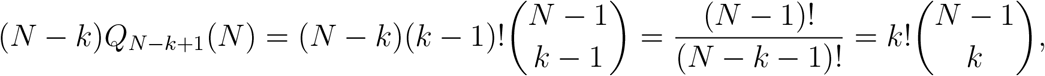

and that, by standard properties of the binomial coefficients, the numerator is

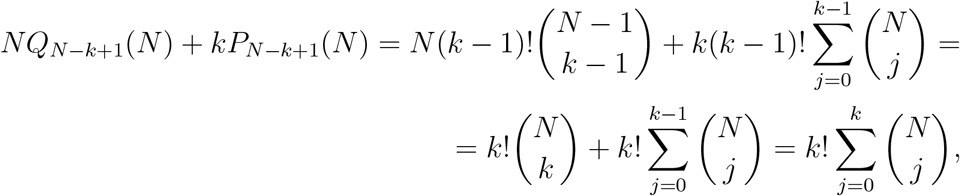

which completes the inductive step and proves the explicit formulas for *P* and *Q*. Lastly, by setting *k* = *N* − 1, we obtain the equation

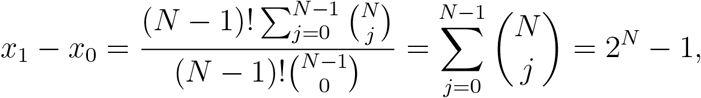

and upon substituting this into the first equation, we obtain the final result

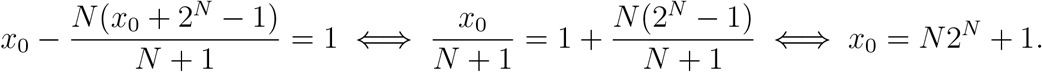

It follows that in this extreme case, the time required to break *σ* grows exponentially with the number of possible mutations *N* .

### The generic case: 0 < μ < 1

We now show that the value of interest, *x*_0_, exhibits exponential, rather than polynomial, growth that when *μ* is bounded away from 1, by repeating the reasoning in the previous section in this regime. More specifically, we show that

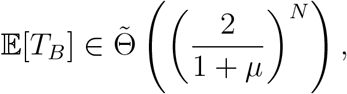

for all 0 *< μ <* 1, where 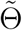 denotes the asymptotic growth order with logarithmic factors neglected.

Because of the more complex structure of the equations involved and the need to extract a single coefficient from the solution of equation (13), we use Cramer’s rule Robinson (1970) to compute 𝔼[*T*_*B*_] as a ratio of two determinants, the denominator being the determinant of the coefficient matrix *I* − *M*_*N*_, and the numerator being the determinant of the same matrix, but with the first column replaced by the right-hand side vector **1**:

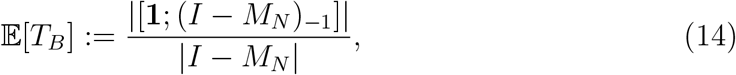

where *X*_−1_ means the matrix *X* without its first column and *X*; *Y* denotes the vertical concatenation of the matrices *X* and *Y* .

### Computing the denominator of (14)

To compute the denominator, we use the standard recurrence used for computing the determinant of a tridiagonal matrix, but start it from the bottom right, rather than the top left. Specifically, if *a*_*i*_, *b*_*i*_, *c*_*i*_ denotes the *i*-th element of the diagonal, sub-diagonal, and super-diagonal, respectively, the recurrence is

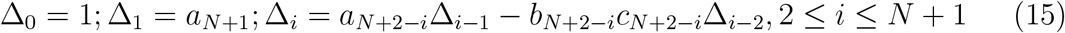

and Δ_*N*+1_ is then the determinant of the matrix (in fact, Δ_*i*_ is the determinant of the matrix consisting of the last *i* rows and columns of the original matrix).

In our case, we have

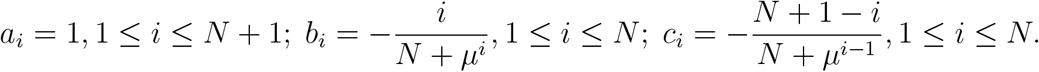

The recurrence then becomes

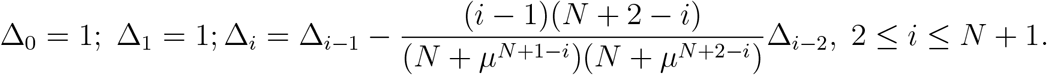

We begin by solving it in the extreme cases *μ* = 0 and *μ* = 1. When *μ* = 0, the recurrence simplifies to

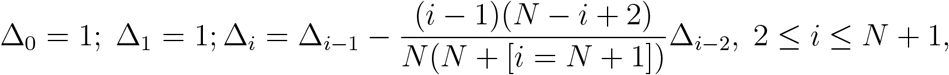

and we will prove by induction that its solution is given by

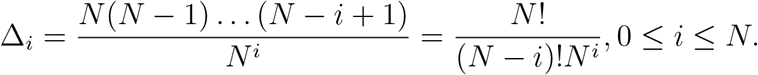

The base cases with *i* = 0 and *i* = 1 correctly produce Δ_0_ = 1 = Δ_1_, and then

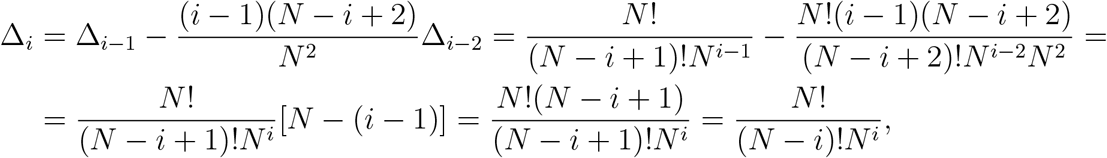

completing the proof by induction.

This means that 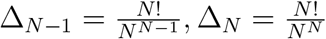 and finally, the recurrence with *i* = *N* + 1 results in 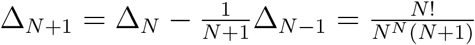. By using Stirling’s approximation, we conclude that, asymptotically for *N* large,

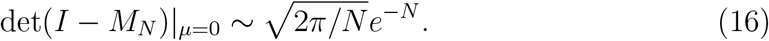

To resolve the case *μ* = 1, we proceed slightly differently; namely, we first scale the matrix *M*_*N*_ by a factor of *N* + 1, and show that the eigenvalues of this scaled matrix, which we denote by *Q*_*N*_, are equally spaced between −*N* and *N* with a step size of 2 (which includes 0 when *N* is even and does not include it when *N* is odd). It will then follow that the eigenvalues of 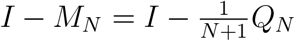 have the form 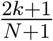 for 0 ≤ *k* ≤ *N*, and the determinant will follow immediately.

Let us consider the scaled matrix *Q*_*N*_ := (*N* + 1)*M*_*N*_, given below.

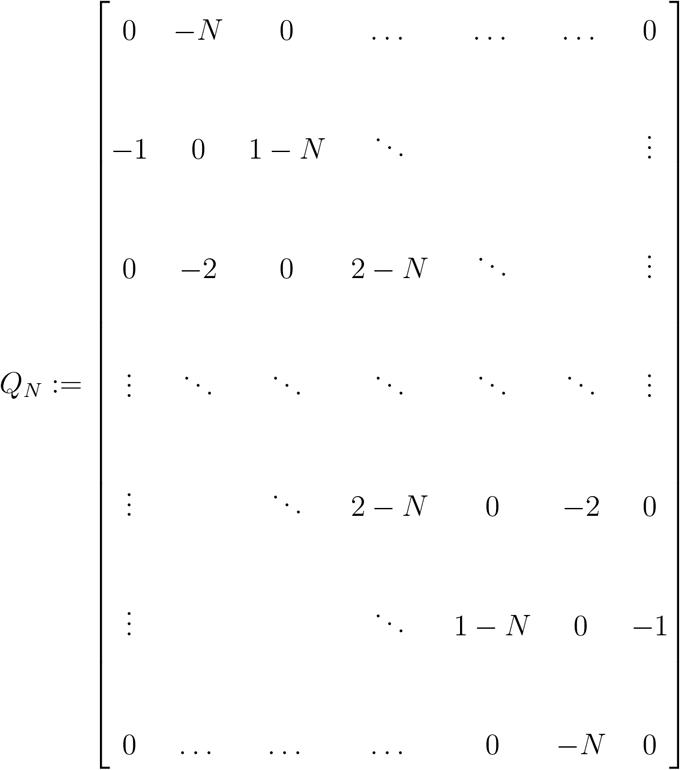

The fact that its eigenvalues are equal to −*N* + 2*k*, 0 ≤ *k* ≤ *N*, is a classical result first published (without proof) by Sylvester in 1854 Sylvester (1854). The first known proof is due to Cayley Cayley (1857), and this matrix was also considered by Schrödinger Schrödinger (1926) who determined the correct eigenvalues, but was unable to prove the result. Here we present the classical treatment due to Taussky and Todd Taussky and Todd (1991), as well as the more sophisticated treatment due to Mark Kac, as reported by Munarini and Torri Munarini and Torri (2005), since it is the latter that we will use to solve the general case.

The elementary approach for factoring the characteristic polynomial of *Q*_*N*_, given (up to sign) by the determinant det(*λI* − *Q*_*N*_), consists of a sequence of steps that do not change its value, followed by a “deflation” procedure that isolates one of the eigenvalues and reduces the dimension by 1. This process is illustrated here for *N* = 3 Taussky and Todd (1991). First, we have

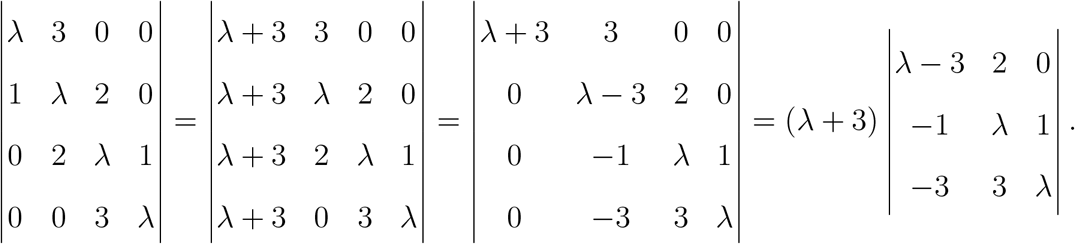

Here, denoting by *R*_*i*_ and *C*_*j*_ the *i*-th row and the *j*-th column, respectively, the first step replaces *C*_1_ with *C*_1_ + *C*_2_ + *C*_3_ + *C*_4_, the second step subtracts *R*_1_ from each of the remaining rows, and the third step expands the determinant along the first column.

Next, we get

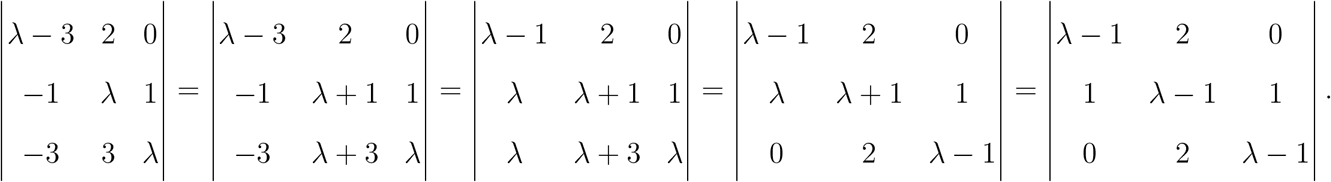

Here, the first step replaces *C*_2_ with *C*_2_ +*C*_3_, the second step replaces *C*_1_ with *C*_1_+*C*_2_, the third step replaces *R*_3_ with *R*_3_ − *R*_2_, and the final step replaces *R*_2_ with *R*_2_ − *R*_1_. This shows that the original matrix *Q*_3_ has −3 as an eigenvalue, and the remaining ones are obtained by adding 1 to those of *Q*_2_, which is easily turned into an inductive proof of the desired statement in the general case.

Since the set 2*k* + 1, 0 ≤ *k* ≤ *N* forms the complete collection of eigenvalues of *Q*_*N*_, it follows that the determinant of *I* − *M*_*N*_ is exactly

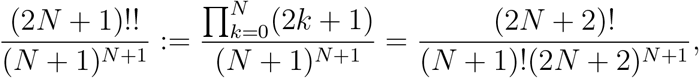

and thus, by Striling’s approximation, asymptotically for large *N*,

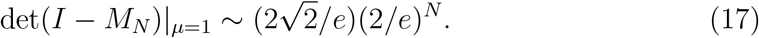

Next, we illustrate the more complex method used by Mark Kac in his approach towards the computation of determinants such as those of *Q*_*N*_ . Towards this end, we define a quantity which is essentially the Cayley continuant Munarini and Torri (2005), but with the signs reversed on the sub- and super-diagonals, which does not change its value as these entries only enter the determinant of a tridiagonal matrix in matching pairs as seen in the recurrence:

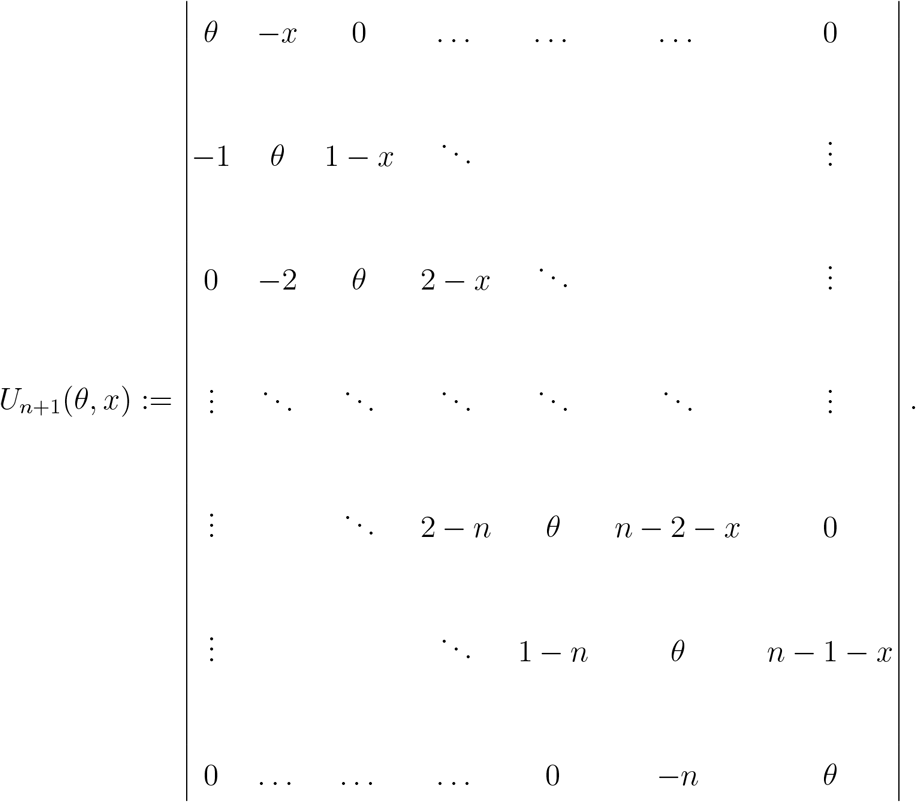

Note that the quantity we defined is indexed by its dimension (hence, the index is *n* + 1 rather than *n*). Clearly, the desired quantity in our case is given by *U*_*N*+1_(*θ* = 0, *x* = *N*).

We increment the index by 1 which adds the entries −(*n* + 1) and (*n* − *x*) at positions (*n* + 2, *n* + 1) and (*n* + 1, *n* + 2), respectively. We then write down the recurrence in (15) in the form

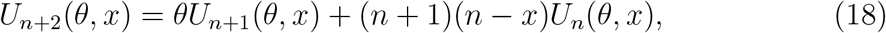

and convert it into a differential equation over the exponential generating series

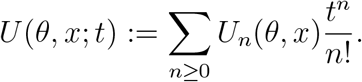

We first multiply each term of (18) by 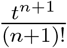 and sum over all *n* ≥ 0. We drop the dependence on *θ* and *x* and use the initial conditions *U*_0_ = 1, *U*_1_ = *θ* to get

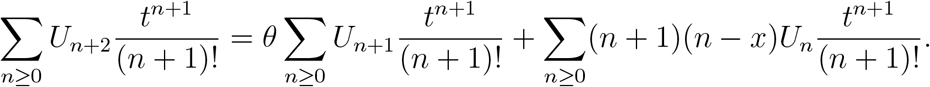

For the left-hand side, note that

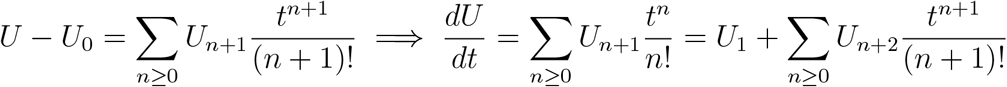

For the right-hand side, the first term simplifies to

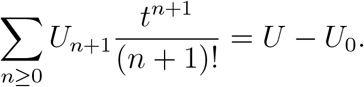

The coefficient of the second term can be written as (*n* + 1)*n* − *x*(*n* + 1), and the first part simplifies to

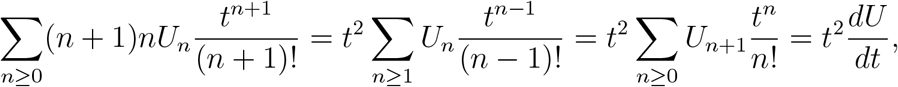

while the second part simplifies to

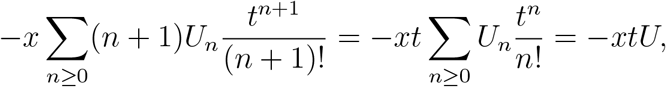

which, in combination with the previous terms, yields the overall result

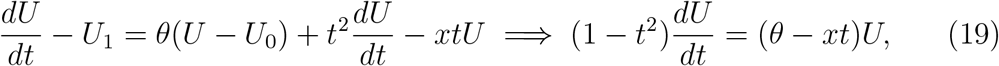

where we used the initial conditions *U*_0_ = 1, *U*_1_ = *θ*. The resulting differential equation is separable, so its solution is readily obtained by standard techniques:

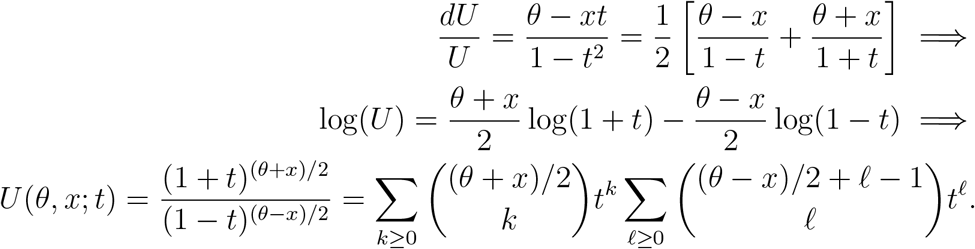

where we use the initial condition *U* (*t* = 0) = *U*_0_ = 1 to identify the integration constant, and the definition of the binomial coefficients as well as the property

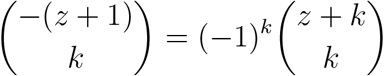

to get the product expression. By expanding the convolution and recalling that *U* is the exponential generating function for the *U*_*n*_(*θ, x*), we finally get

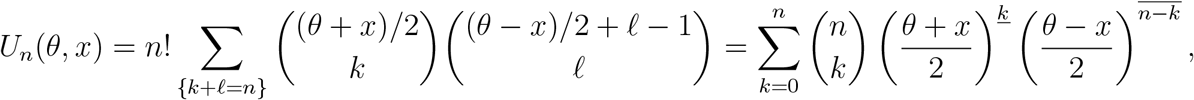

where the falling factorial 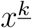 and rising factorial 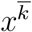 are respectively defined by

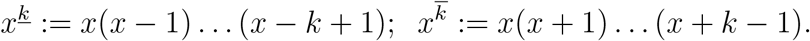

In particular, this implies the desired result

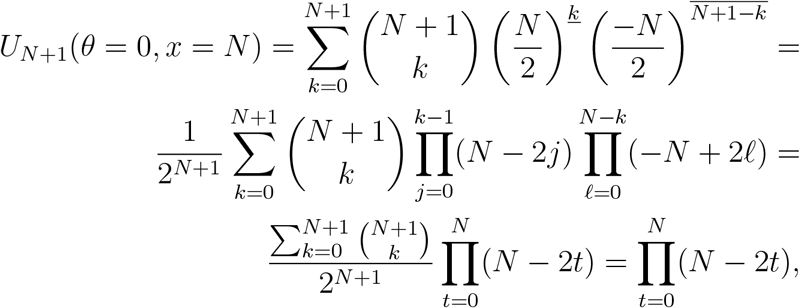

in agreement with our earlier result.

We will now proceed with computing the determinant in the general case. We will proceed similarly to the above, but alter the diagonal to contain powers of *μ* in addition to the constant, defining *V*_*n*_ as the resulting determinant:

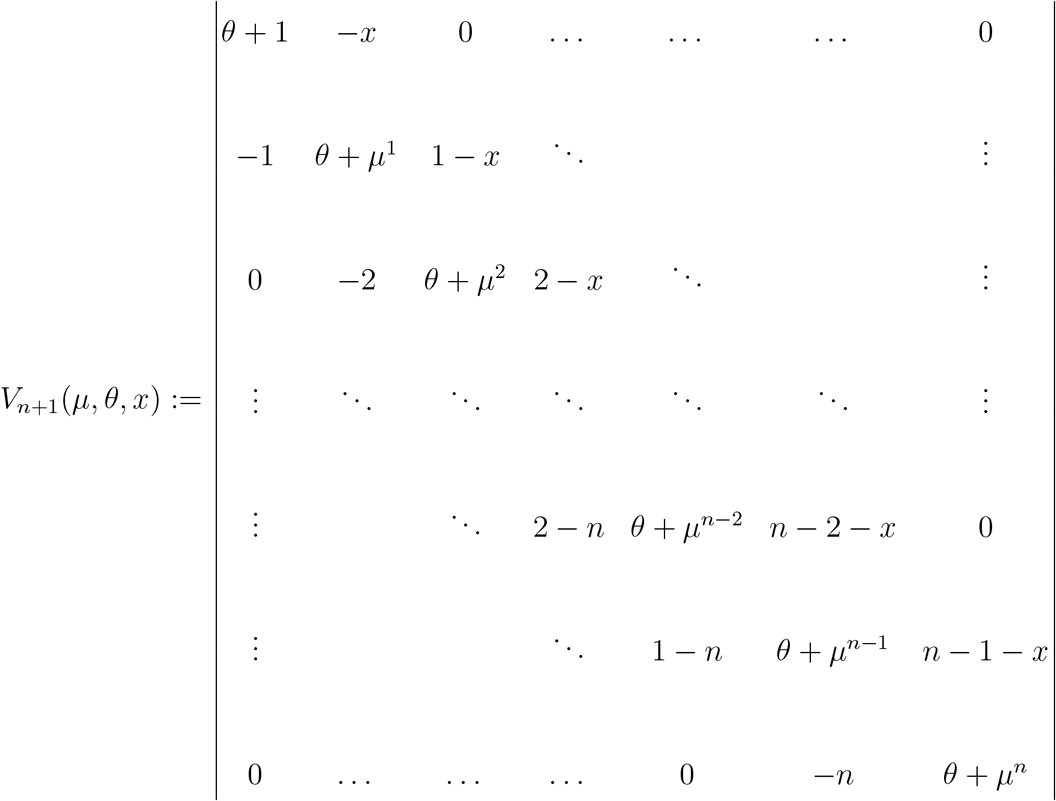

and the value of interest to us is *V*_*N*+1_(*μ* = *μ, θ* = *N, x* = *N*)*/D*_*N*+1_(*μ*), where

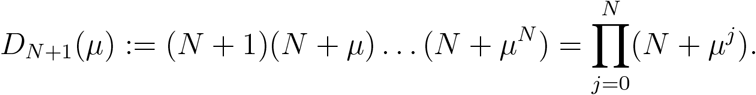

It is immediate that *D*_*N*+1_(*μ*) is an increasing function of *μ* with *D*_*N*+1_(*μ* = 0) = *N*^*N*^ (*N* + 1) and *D*_*N*+1_(*μ* = 1) = (*N* + 1)^*N*+1^, and it follows that in general, for all ≤ *μ* ≤ 1, we have *D*_*N*+1_(*μ*) = *c*(*N, μ*)*N*^*N*^ (*N* + 1), with

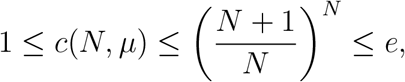

so *D*_*N*+1_(*μ*) may be treated as independent of *μ* up to a multiplicative constant.

Rather than solving for *V*_*N*+1_ via a differential equation (which becomes a functional-differential equation due to the presence of *μ*) we simplify our task by expanding it as a polynomial in powers of *μ* and truncating the result.

Since *V*_*N*+1_ as a function of *μ* is a polynomial of degree *N* (*N* + 1)*/*2, let us write 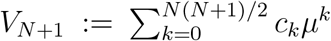. Let *M* be the matrix used to define *V*_*N*+1_. Since the determinant of *M* is obtained as a (signed) sum of *M*_*i,π*(*i*)_ over all permutations *π* of *N* + 1 elements, we can also write this differently by grouping the permutations according to the set of diagonal terms (the fixed points of *π*) from which we select powers of *μ* (as opposed to *N*). If *S* ⊆ [*N* + 1] is this subset, the remaining products in aggregate equal the determinant of the matrix *M*_−*S*,−*S*_ obtained by crossing out the rows and columns corresponding to *S* and setting *μ* = 0 in what is left (here, we denote by [*L*] the set of all integers between 1 and *L* inclusively). In other words, defining |*S*| := ∑_*s*∈*S*_ *s*,

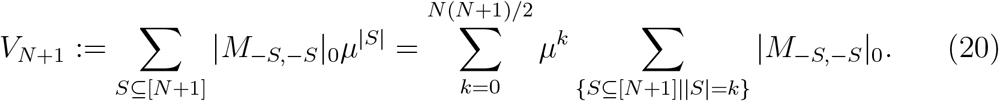

In particular, note that the coefficient of *μ*^0^ in *V*_*N*+1_ is obtained by using the expression for Δ_*N*+1_ when *μ* = 0 and multiplying it by *N*^*N*^ (*N* + 1), the scaling factor. It follows that this coefficient is given by

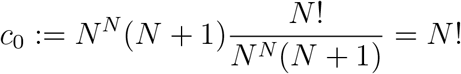

We make two observations about the remaining coefficients *c*_*k*_. First, the terms corresponding to *μ*^*k*^ with *k > N* do not make a major contribution to the total because they require sets *S* of size at least 2, and the corresponding terms involve selecting several times the powers of *μ* from the diagonal terms, which are much smaller than the other component of the diagonal term, *N*, since 0 *< μ <* 1. Although there are many such sets, we will see that their contribution can be neglected.

Second, the coefficients corresponding to *μ*^*k*^ with 1 ≤ *k* ≤ *N* are dominated by the term corresponding to the singleton *S* = {*k*}, by the same reasoning. We will therefore obtain *c*_*k*_ to leading order with respect to *N* by computing the (*k, k*) co-factor of *V*_*N*+1_, i.e. the determinant of the matrix obtained by crossing out its *k*-th row and column, with *μ* = 0.

It is easy to see that this cofactor splits into the product of two terms, say *c*_*k*_ = *d*_*k*_*e*_*k*_. The first of these, *d*_*k*_, is the determinant of the submatrix formed by the first *k* rows and columns of the matrix defining *V*_*N*+1_, and the second one, *e*_*k*_ (up to a scaling factor of *N*^*N*−*k*^), equals Δ_*N*−*k*_ when *μ* = 0, i.e. the determinant of the submatrix formed by the last *N* − *k* rows and columns of *M*_*N*_ when *μ* = 0, computed previously, so

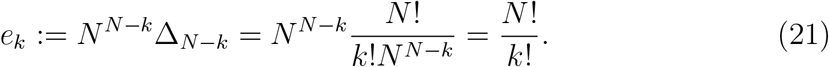

It remains to compute *d*_*k*_, obtained as *V*_*k*_ evaluated at *μ* = 0, *θ* = *x* = *N* . This can be achieved by altering the initial conditions in the original equation for *U*, (18), to *U*_0_ = 1, *U*_1_ = *θ* + 1. After an identical transformation this results in the inhomogeneous equation

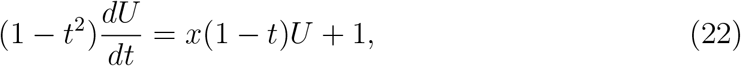

where we use the fact that *θ* = *x* = *N* in our case of interest here. Using the homogeneous solution as an ansatz, we set *U* (*t*) = (1 + *t*)^*x*^*f* (*t*) and this results in a simpler differential equation for *f* (*t*),

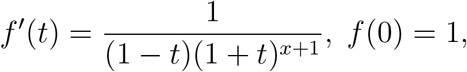

whose solution is obtained by a partial fraction expansion of the right-hand side:

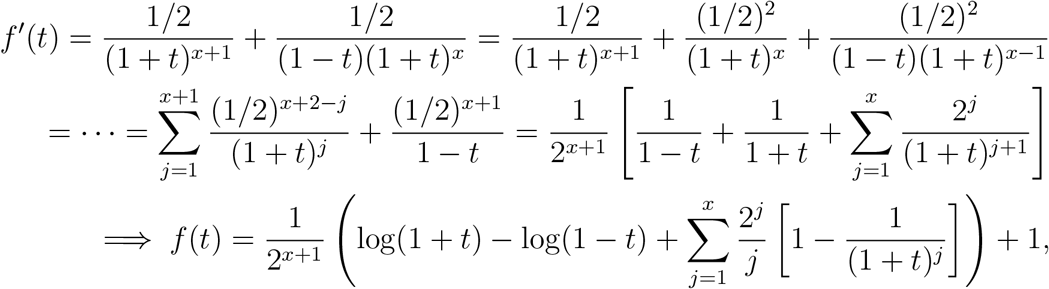

where the last term and the square brackets account for the integration constant.

The expression *U* (*t*) = (1 + *t*)^*x*^*f* (*t*) with *x* = *N* finally becomes

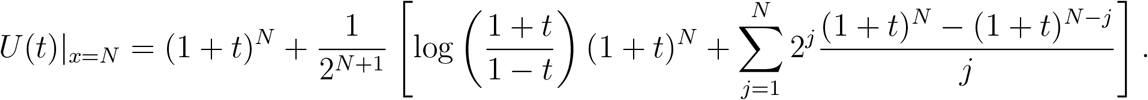

We note the following fact: since 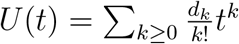 and 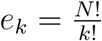, we have

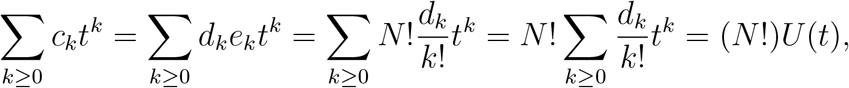

and therefore, the expression of interest equals (*N* !)*U*_≤*N*_ (*t* = *μ*), where *U*_≤*N*_ (*t*) denotes the truncated series

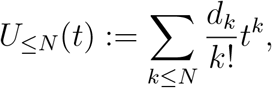

noting that the two expressions agree at 0 since *c*_0_ = *N* ! (as shown previously).

To estimate this truncated series we note that the only part of *U* (*t*) whose expansion extends beyond *t*^*N*^ is log(1 + *t*) − log(1 − *t*) (1 + *t*)^*N*^ . We compute the series expansion of log(1 + *t*) − log(1 − *t*), which is given by 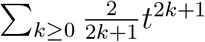. Since all the coefficients of both log(1 + *t*) − log(1 − *t*) and (1 + *t*)^*N*^ are non-negative, for 0 ≤ *t* ≤ 1 we have

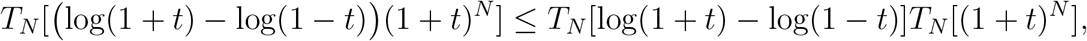

where *T*_*N*_ is the truncation operator on a power series that returns its first *N* + 1 terms; this holds for the convolution product of any two power series with non-negative coefficients since

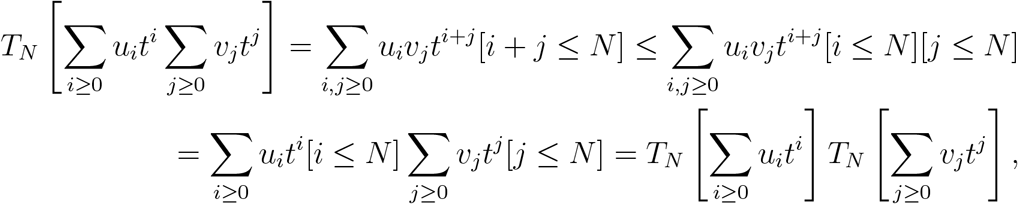

where [*E*] is the Iverson bracket whose value is 1 is expression *E* is true and 0 otherwise.

We also see that, on 0 ≤ *t* ≤ 1, since 2(*N* − 1) + 1 ≥ *N* for all *N* ∈ ℕ, we get

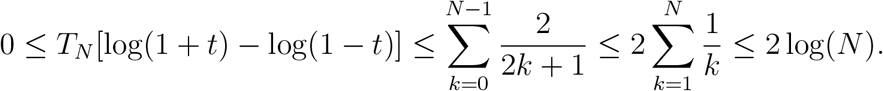

It follows that

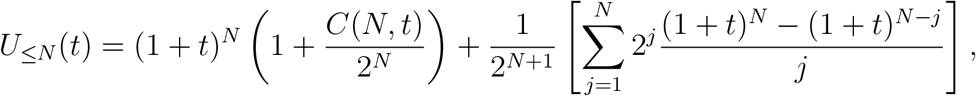

where 0 ≤ *C*(*N, t*) ≤ log(*N*) uniformly for all *N* and 0 ≤ *t* ≤ 1.

To estimate the second term of *U*_≤*N*_ (*t*), we let 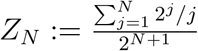 and compute

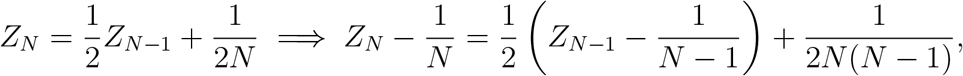

and since *Z*_*N*_ is bounded above by 1 and below by 1*/N*, it follows that *Z*_*N*_ − 1*/N* must converge to 0 as *N* increases. It therefore follows that

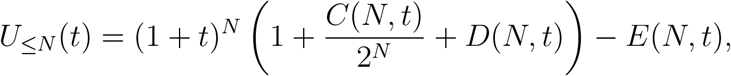

where *D*(*N, t*) ≥ 0, *D*(*N, t*) − 1*/N* → 0 as *N* → ∞ and 0 ≤ *E*(*N, t*) ≤ log(*N*) as

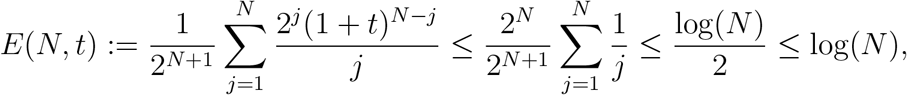

uniformly for all *N* and 0 ≤ *t* ≤ 1, so that finally, *U*_≤*N*_ (*t*) ≈ (1 + 1*/N*)(1 + *t*)^*N*^ .

When we compute (*N* !)*U*_≤*N*_ (*t* = 1), using this approximation results in 2^*N*^ (*N* + 1)!*/N*, giving the denominator of (14) as 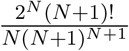, which Stirling’s formula shows to be in agreement with the previously found result, equation (17), up to a factor of 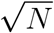, which is therefore the cost of approximating the polynomial in (20) using only sets of size at most 1.

### Computing the numerator of (14)

To compute the numerator of (14) we need to evaluate

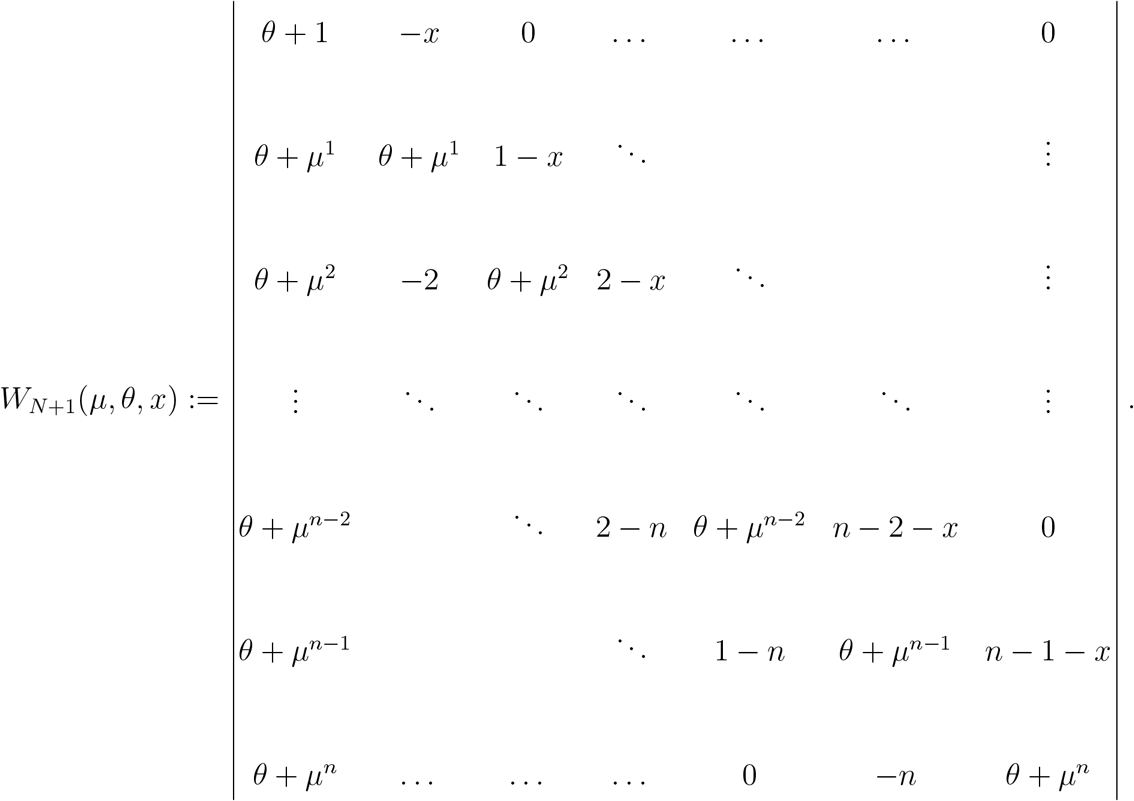

The value of interest to us will then be *W*_*N*+1_(*μ* = *μ, θ* = *N, x* = *N*)*/D*_*N*+1_(*μ*), where *D*_*N*+1_(*μ*) remains exactly as defined in the previous subsection.

To evaluate *W*_*N*+1_ we begin by expanding it along the first column to get

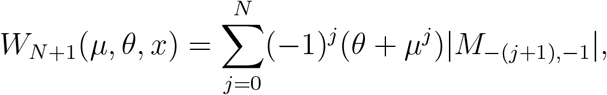

where *M* is the matrix whose determinant is *W*_*N*+1_. Furthermore, we note that because *M* is tridiagonal, all the terms with the exception of the one with *j* = 0 split into a product of two terms, the first of which is the product of the first *j* − 1 super-diagonal entries and the second of which is the determinant of the last *N* − *j* rows and columns of *M* . More precisely:

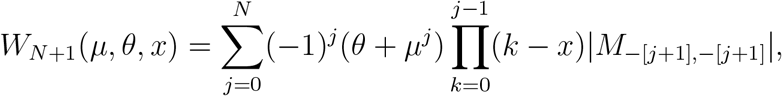

where, as before, [*L*] denotes the set of all integers from 1 to *L*.

This is because the shape of the matrix remaining after crossing out the *j* + 1-th row (using the numbering from 1 to *N* + 1) and the first column of *M* is a tridiagonal matrix with non-zero entries on the diagonal, sub-diagonal and sub-sub-diagonal (rather than the super-diagonal) above the original *j* + 1-th row, and a “standard” tridiagonal matrix below it.

The critical entry to consider is the entry *M*_*j*+2,*j*+1_ in the original matrix, which becomes entry (*j* + 1, *j*) in the truncated matrix. We show that any permutation *π* ∈ *S*_*N*_ that contains this entry (i.e. that has *π*(*j* + 1) = *j*) produces a zero product for structural reasons, and does not contribute to the determinant. Indeed, crossing out the *j* + 1-th row and the *j*-th column in the truncated matrix leaves an imbalance - the top left corner is a possibly empty *j ×* (*j* − 1) submatrix, the bottom right corner is a possibly empty (*N* − *j* − 1) *×* (*N* − *j*) submatrix, and the bottom left and top right corners both only contain 0’s, meaning that any permutation yields a zero product, as claimed.

It now follows that the determinant of the truncated matrix factors into the product of two determinants. The first one is the determinant of the *j × j* tridiagonal matrix with non-zero entries only on and below the diagonal, which is therefore also lower-triangular and thus, its determinant equals the product of its diagonal entries, which were the first *j* super-diagonal entries of *M* . The second one is the determinant of the *N* − *j × N* − *j* tridiagonal matrix containing the last *N* − *j* rows and columns of *M*, leading to the desired conclusion.

We now recall that, by the reasoning in the previous section, the dominant contributions to various terms involving *μ* in the definition of *W*_*N*+1_ come from permutations where *μ*^*k*^ is a monomial arising from the selection of *μ*^*k*^ in the *k, k*-th entry of *M* while the remainder is approximated by setting *μ* = 0 in the corresponding determinant. It follows that

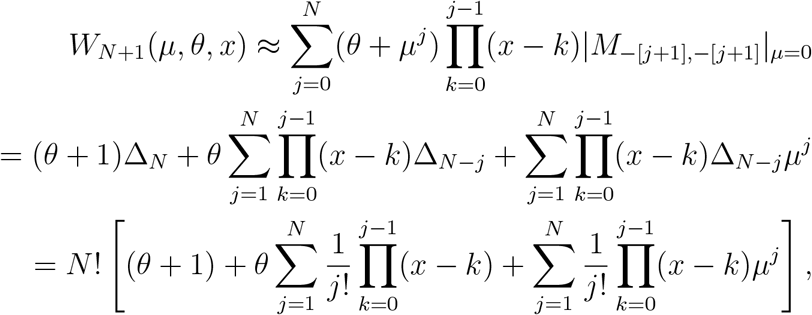

where we distributed the negative sign across the product in the first line and used the definitions *e*_*j*_ = *N*^*N*−*j*^Δ_*N*−*j*_ = |*M*_−[*j*+1],−[*j*+1]_|_*μ*=0_ in the second line and their known values from (21) in the third line.

Upon substituting *θ* = *x* = *N* in the above, we finally obtain

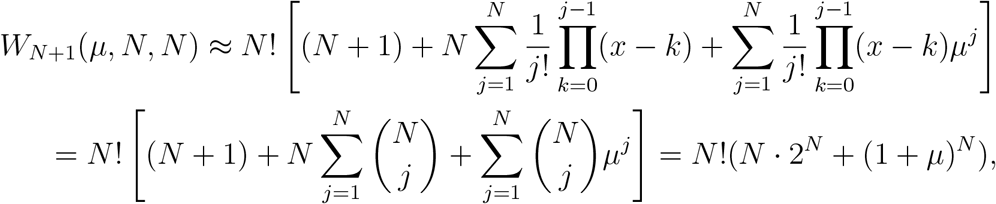

where we used the Binomial theorem in the last line. We verify that this yields correctly yields *N* !(*N* · 2^*N*^ + 1) when *μ* = 0 and (*N* + 1)! · 2^*N*^ when *μ* = 1, the latter value being once again within a factor of 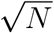 from the correct one, (2*N* + 1)!! · (*N* + 1), according to Stirling’s theorem and the work in equation (17).

The overall conclusion is that, asymptotically for *N* large and 0 ≤ *μ* ≤ 1, the expected time scales as

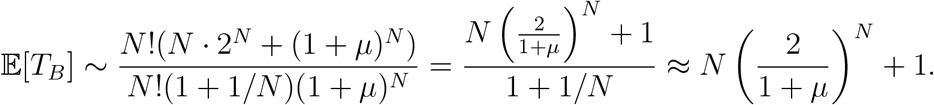

Note that, although the right-hand side happens to be exact for both *μ* = 0 and *μ* = 1, it may be off by as much as a factor of *N* in either direction for values strictly between 0 and 1 due to approximation errors; however, the rate of growth remains exponential in *N* provided that *μ* is bounded away from 1. However, numerical experiments suggest that the true value tends to be much closer to this estimate in practice, with relative errors on the order of 1*/N*, meaning that in fact, the errors in the approximation of the numerator and of the denominator may be in the same direction.

### Details to make the proof rigorous

In this section we include additional details required to make the proof rigorous. The main result is the following

#### Lemma 0.1.

*Let* 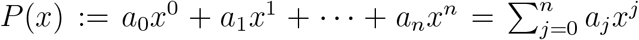 *be a polynomial with all a*_*i*_ *non-negative and a*_0_ *>* 0. *Let* 1 *< k* ≤ *n be given and let* 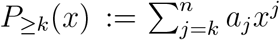 *be the truncated version of this polynomial. Then, for any t >* 0, *the relative truncation error P*_≥*k*_(*x*)*/P* (*x*) *is non-decreasing on* [0, *t*].

*Proof*. Let 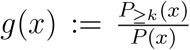 be the truncation ratio. Suppose, on the contrary, that there exist 0 ≤ *t < t*′ such that *g*(*t*) *< g*(*t*′). Equivalently, denoting the sum of the first *k* terms of *P* (*x*) by *P*_*<k*_(*x*), and using the non-negativity of *P* (*x*) and of *P*_*<k*_(*x*) for *x >* 0, this means that

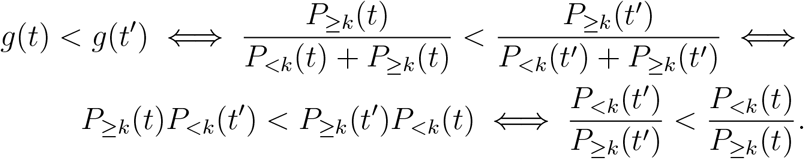

Now the ratio *f* (*x*) := *P*_≥*k*_(*x*)*/P*_*<k*_(*x*) is a rational function whose denominator does not vanish for *x* ≥ 0, and is therefore continuously differentiable on [0, *t*]. Therefore, by Fermat’s theorem, it can only attain an extremum inside [0, *t*] at a stationary point, where its first derivative vanishes. However, we have

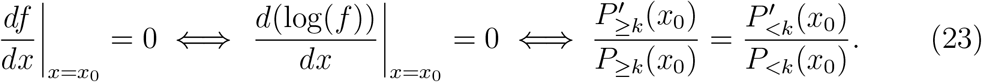

Note that, given *L* non-negative real numbers *w*_1_, *w*_2_, …, *w*_*L*_, *L* non-negative real numbers *p*_1_, *p*_2_, …, *p*_*L*_, and *L* positive real numbers *q*_1_, *q*_2_, …, *q*_*L*_, we have

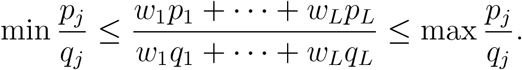

Applying the right-hand inequality to the left-hand side of the equality in equation (23), with *L* = *k* and *w*_*j*_ = *a*_*j*−1_, and assuming that *x*_0_ *>* 0, we get

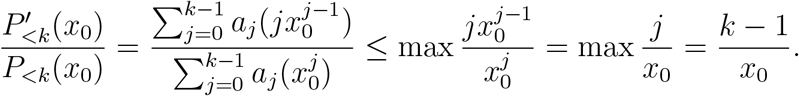

On the other hand, applying the left-hand inequality to the right-hand side of the equality in equation (23), with *L* = *n* + 1 − *k* and *w*_*j*_ = *a*_*k*+*j*_, we get

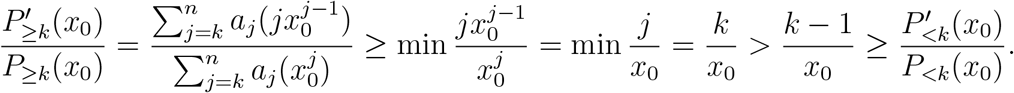

It follows that the equality in equation (23) cannot be attained inside [0, *t*], so the derivative of *f*′(*x*) cannot change sign on [0, *t*]. Since *f* (*x*) is positive near 0, it remains positive on [0, *t*] and hence, *f* (*x*) is non-decreasing on [0, *t*], so 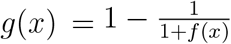 is also non-decreasing on [0, *t*], completing the proof. □

## Notes

### Competing Interest Statement

The authors have declared no competing interest.

### Summary of Updates

Updated author name (there was a typo)

